# Early invasion of uropathogenic *Escherichia coli* into the bladder wall by solitary bacteria that are protected from antibiotics and neutrophil swarms in an organoid model

**DOI:** 10.1101/2020.10.29.358622

**Authors:** Kunal Sharma, Vivek V. Thacker, Neeraj Dhar, François Signorino-Gelo, Maria Clapés Cabrer, Anaëlle Dubois, Jasper Mullenders, Graham Knott, Hans Clevers, John D. McKinney

## Abstract

Uropathogenic *Escherichia coli* (UPEC) is the most common cause of urinary tract infections (UTIs) requiring antibiotic therapy. Recurrent infections, which occur in a quarter of treated individuals, may arise from “quiescent intracellular reservoirs” of bacteria that invade deeper layers of the bladder wall following infection and exfoliation of superficial umbrella cells. Here, we present a novel bladder organoid model of UPEC infection that recapitulates the stratified bladder architecture within a small volume suitable for live-cell imaging of host-pathogen dynamics with high spatiotemporal resolution. We confirm that bacteria injected into the organoid lumen rapidly enter superficial cells that resemble umbrella cells and proliferate to generate tightly packed colonies that resemble intracellular bacterial communities (IBCs), a hallmark of UPEC pathogenesis. Unexpectedly, at early stages of infection we detect individual “solitary” bacteria that penetrate deeper layers of the organoid wall, where they evade killing by antibiotics and neutrophils. Volumetric serial block face scanning electron microscopy of infected organoids reveals that solitary bacteria can be found throughout the bladder wall and may be intracellular or pericellular (sandwiched between uroepithelial cells). Unlike bacteria within IBCs, which are coccoid-shaped and non-flagellated, solitary bacteria within the bladder wall are rod-shaped and flagellated. We conclude that early invasion of deeper layers of the bladder wall, independent of IBC formation, results in the establishment of reservoirs of solitary bacteria that resist elimination by antibiotics and the host innate immune response.

## Introduction

Urinary tract infections (UTIs) are among the most common bacterial infections and the second most-common cause for the prescription of antibiotics (Foxman, 2010). Although seldom fatal, UTIs substantially reduce the quality of life and incur enormous healthcare costs. Recurrence, defined as a reappearance of infection within 12 months despite the apparently successful completion of antibiotic therapy, occurs in a quarter of all UTIs (Foxman et al., 2000). Women are at particularly high risk, and more than 60% of women will be diagnosed with a UTI at least once in their lifetime (Klein and Hultgren, 2020). Uropathogenic *Escherichia coli* (UPEC) is the causative agent of about 80% of all UTIs, which can be further complicated by dissemination of UPEC from the bladder (cystitis) to the kidneys (nephritis) or into the bloodstream (urosepsis).

Much of our current understanding of UPEC pathogenesis has been derived from experiments in mouse models of infection(Hannan and Hunstad, 2016; Hung et al., 2009), which reveal that UPEC can grow extracellularly in the urine within the bladder (Alteri et al., 2009; Forsyth et al., 2018; Hull and Hull, 1997; Subashchandrabose et al., 2014) or intracellularly in the bladder wall (Anderson et al., 2003; Justice et al., 2004; Mulvey et al., 2001). Invasion into the bladder wall is preceded by adherence of UPEC to the superficial umbrella cell layer of the bladder, mediated by interactions of the bacterial type I pilus with uroplakin proteins on the surface of the uroepithelium (Klein and Hultgren, 2020; Lewis et al., 2016; Martinez et al., 2000; Mulvey et al., 1998). Following bacterial entry into umbrella cells, a subset of bacteria proliferate to form biofilm-like “intracellular bacterial communities” (IBCs) (Anderson et al., 2003; Justice et al., 2004; Mulvey et al., 2000) which protect the bacteria from clearance by antibiotic treatment (Blango and Mulvey, 2010) or innate immune responses (Justice et al., 2004). At later stages of infection, UPEC penetrates into deeper layers of the bladder to form “quiescent intracellular reservoirs” that may also be responsible for recurrent infection after antibiotic therapy (Mulvey et al., 2000, 2001; Mysorekar and Hultgren, 2006; Schilling et al., 2002). However, these isolated subpopulations have been difficult to characterize in bladder sections, and the relative contribution of IBCs and quiescent intracellular reservoirs in recurrent infections remains unclear (Blango et al., 2014; Mulvey et al., 2001; Scott et al., 2015).

The number of bacteria in three identifiable subpopulations (extracellular bacteria, IBCs, and quiescent intracellular reservoirs), their relative growth dynamics, and their survival under attack from antibiotics or immune cells are difficult to characterize *in situ* in animal models. Many studies have therefore relied on an examination of bladder explants at specified time points post-infection. Although powerful, this technique does not provide information on the underlying dynamics of host-pathogen interactions, nor does it permit the quantification of *in situ* growth of extracellular bacteria (Anderson et al., 2003; Justice et al., 2004). *In vitro* models have been developed to study specific aspects of UPEC infection, such as the role of the stratified bladder architecture (Horsley et al., 2018) or the effects of micturition on IBC formation (Andersen et al., 2012). However, these models suffer from limitations in their ability to recreate a stratified epithelium with multiple differentiated cell layers (Andersen et al., 2012), and three-dimensional migration of immune cells into the bladder in response to infection is difficult to model in platforms based on Transwell inserts (Horsley et al., 2018). In many of these systems, live-cell imaging remains technically challenging (Horsley et al., 2018; Smith et al., 2006).

In the last decade, organoids have emerged as experimentally tractable biomimetic systems that recapitulate key physiological and functional features of the cognate organs (Clevers, 2016). These complex 3D multicellular structures are generated from stem cells or organ-specific progenitor cells (Rossi et al., 2018) and have now been established for a number of different organs (Liu et al., 2004; Sachs et al., 2019), including the bladder (Lee et al., 2018; Mullenders et al., 2019). Recently, organoids have also emerged as model systems to study host-pathogen interactions during infections caused by bacteria (Bartfeld and Clevers, 2015; Co et al., 2019; Kessler et al., 2019; Pleguezuelos-Manzano et al., 2020; Williamson et al., 2018), viruses (Qian et al., 2017; Sanden et al., 2018; Zhou et al., 2018), or parasites (Heo et al., 2018; Nikolaev et al., 2020).

Bladder organoids offer several distinct advantages as model systems for UTIs. They recapitulate the stratified and differentiated layers of the uroepithelium, possess a central lumen that mimics the bladder lumen, and are easier to manipulate than whole-animal infection models. Importantly, the compact volume of individual organoids can be imaged in its entirety with high spatiotemporal resolution using time-lapse confocal microscopy. This makes it possible to follow the rapidly changing dynamics of UTIs and to monitor the responses of host cells and bacterial cell to external perturbations, such as a defined course of antibiotic treatment or the addition of innate immune cells (Neal et al., 2018; Sachs et al., 2019). Compared to conventional *ex vivo* tissue explants (Justice et al., 2004) or *in vitro* infection models (Andersen et al., 2012) bladder organoids offer a more realistic reconstitution of bladder physiology that is accessible to a wider range of experimental techniques for studies of UPEC pathogenesis.

Here, we establish a model system to study the early dynamics of UPEC pathogenesis based on mouse bladder organoids derived from primary cells (Mullenders et al., 2019). We use a combination of time-lapse laser scanning confocal microscopy and serial block face scanning electron microscopy (SBEM) to monitor the dynamics of UPEC invasion, growth, and persistence within bladder organoids with single-cell and sub-micron resolution. We find that individual “solitary” bacteria are seeded in deeper layers of the bladder epithelium concomitantly with but independently of the formation of IBCs in the superficial layer of umbrella-like cells. These solitary bacteria are refractory to killing by antibiotics and neutrophils, suggesting that early invasion of bacteria into deeper layers of the bladder wall may play an important role in recurrent infections.

## Results

### Establishment of differentiated mouse bladder organoids

We generated mouse bladder organoids from C57BL/6 wild-type (WT) or mT/mG mice (Muzumdar et al., 2007), which express the red-fluorescent protein tdTomato within cell membranes, following a recently published procedure (Fig. 1A) (Mullenders et al., 2019). Briefly, mouse bladders were isolated *ex vivo* and epithelial cells extracted from the bladder lumen were cultured in basement membrane extract to generate organoids over a period of 2-3 weeks (Fig. 1A). We verified that these organoids recapitulate the different layers of the stratified mouse uroepithelium by comparative immunofluorescence staining of bladder organoids and explanted mouse bladders. Staining slices of organoids with antibodies directed against uroplakin-3a (UP3a) or cytokeratin 8 (CK8) showed that cells lining the lumen of the bladder organoids highly expressed both markers (Fig. 1B). This staining pattern is consistent with that observed in the umbrella-cell layer in slices of mouse bladder tissue (Fig. 1C). However, we could not rule out the possibility that some of the UP3a+ cells in the organoid might be underlying transitional bladder cells (Mysorekar and Hultgren, 2006). The presence of intermediate and basal cell layers in the bladder organoids was confirmed by staining with antibodies directed against cytokeratin 13 (CK13) (Fig. 1B,), p63 (Fig. S1A,), or cytokeratin 7 (CK7) (Fig. S1B,), which were also consistent with the staining patterns observed in mouse bladder slices (Fig. 1C, S1C, S1D). In addition, CK8 expression was higher in the umbrella-like cell layer (demarcated by a white dashed line in Fig. 1B) compared to the underlying intermediate and basal layers in both the organoids and mouse bladder tissue (Southgate et al., 1994). A 3D view of an entire bladder organoid stained with antibodies against CK8 confirmed that the cell layer surrounding the lumen in the centre of the organoid is CK8+ (Fig. S2). Bladder organoids therefore recapitulate the stratified architecture of the mouse uroepithelium with high levels of uroplakin expression in the superficial umbrella-like cell layer, which is important for UPEC adherence and invasion mediated by type I pili (Mulvey et al., 1998).

**Figure 1:**
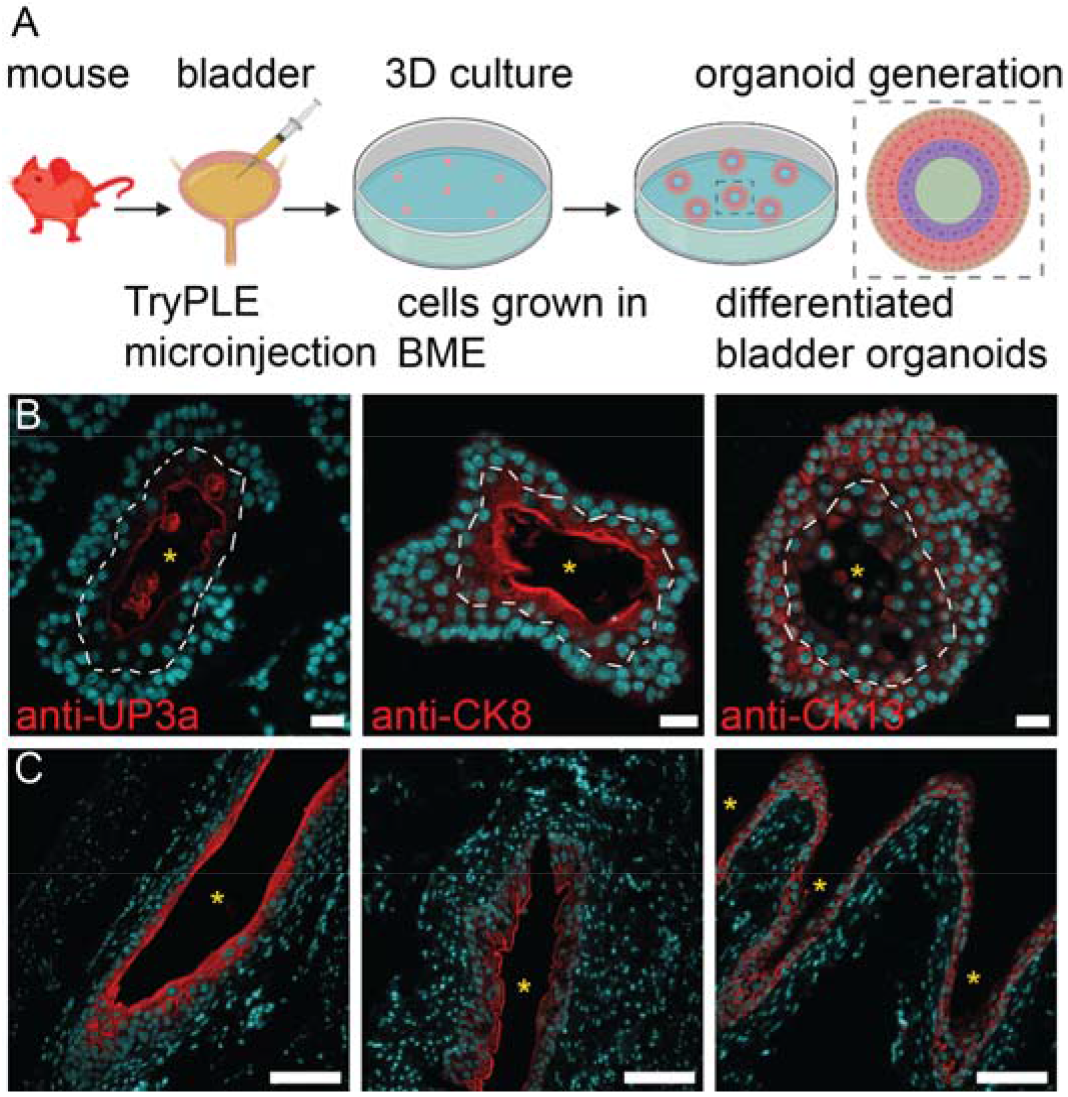
Mouse bladder organoids recapitulate the stratification of bladder uroepithelium. (**A**) Schematic of the protocol used for generation of mouse bladder organoids. Lumenal cells, isolated by microinjection of TryPLE solution into the bladder, are cultured in basement membrane extract (BME) to form differentiated bladder organoids. (**B, C**) Immunofluorescence staining confirms that bladder organoids (**B**) recapitulate the stratified layers of the uroepithelium observed in mouse bladder tissue (**C**). The umbrella-cell layer was identified with anti-uroplakin 3a (anti-UP3a) and anti-cytokeratin 8 (anti-CK8) antibodies. The basal and intermediate cell layers were identified with anti-cytokeratin 13 (anti-CK13) antibody. Cell nuclei were labeled with DAPI (cyan). The lumens of the bladder organoids and mouse bladder slices are indicated by yellow asterisks. The dashed white lines demarcate the boundary between the umbrella-like cell and intermediate cell layers. Scale bars, 20 μm in (**B**) and 100 μm in (**C**).

### Early invasion of UPEC into the bladder wall provides protection against antibiotics

Intravital imaging of antibiotic treatment and recovery in the infected bladder is extremely challenging(Justice et al., 2004, 2006). We therefore modelled the acute phase of a UTI by microinjecting UPEC expressing yellow fluorescent protein (YFP) into the lumen of individual bladder organoids, which mimics the natural route of infection through the urethra (Fig. 2A). The infection cycle was modelled in three stages: stage one (0-165 minutes), an initial period of unimpeded bacterial proliferation; stage two (165-345 minutes), treatment with ampicillin at ten-fold the minimum inhibitory concentration (10X-MIC, corresponding to 64.5 μg/mL); stage three (345-525 minutes), bacterial recovery after ampicillin washout. Snapshots from time-lapse imaging of two infected organoids are shown in Fig. 2B-E (see supplementary movie SMov1) and Fig. 2F-I (see supplementary movie SMov2); an additional example is shown in Fig. S3A1-A7 (see supplementary movie SMov3). We microinjected organoids with between 300 and 1300 colony forming units of UPEC bacteria in a 1nl volume (inocula in Table S1), although the number of bacteria that are retained in the organoid after the subsequent washing steps is about 10-fold lower. Soon after microinjection, rapid bacterial growth within the organoid is observed (Fig. 2B, C and 2F, G), predominantly within the lumen (indicated by a yellow arrowhead in Fig. 2B). Bacterial growth is measured using raw Z-stacks processed via an analysis pipeline in Imaris Bitplane (see Materials and Methods and Fig. S4). Briefly, the fluorescence intensity of tdTomato in uroepithelial cells and YFP in bacteria is used to segment organoid and bacterial volumes, which are depicted as surfaces whose properties are measured using standard algorithms in Imaris.

**Figure 2.**
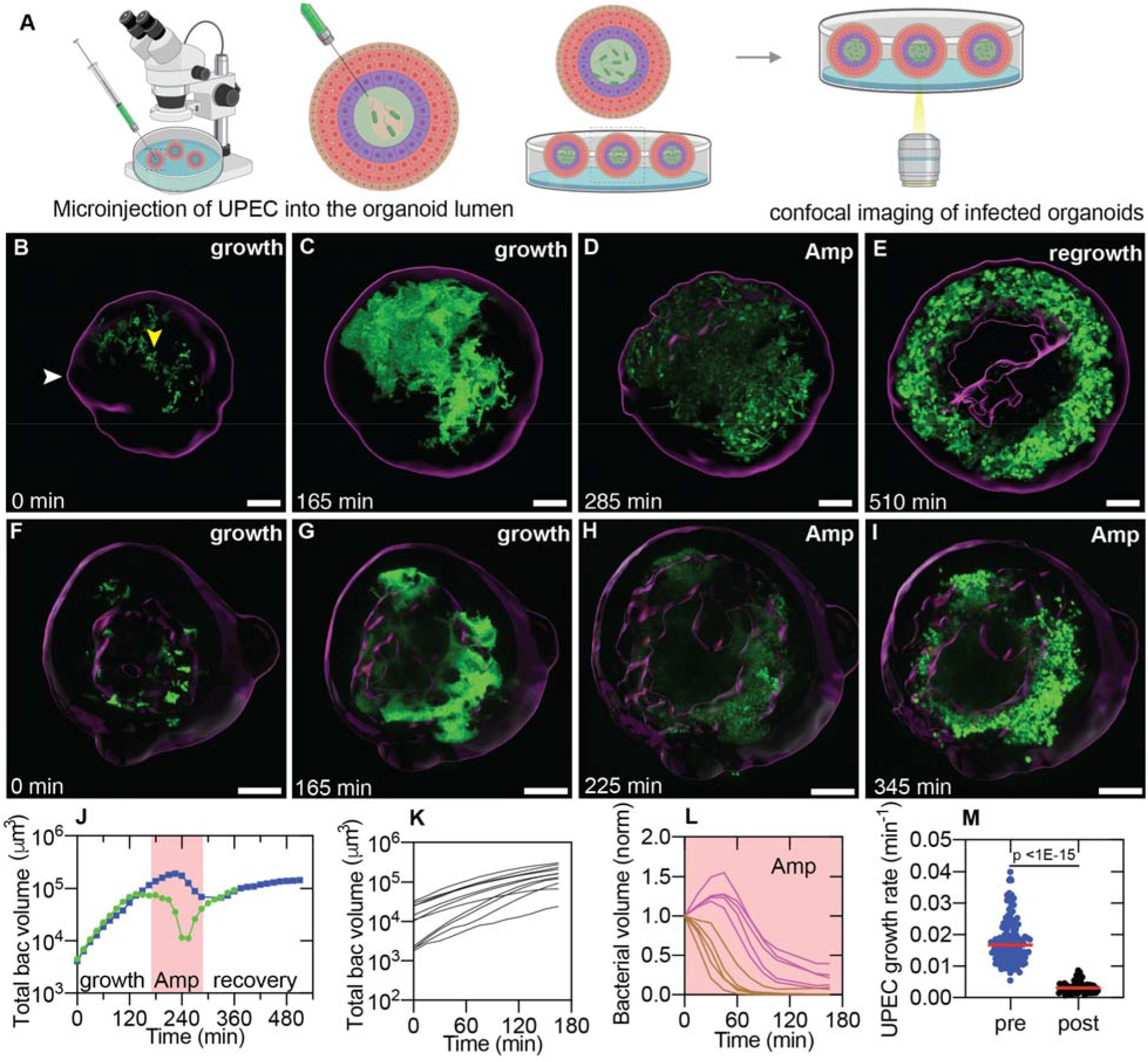
Bacteria within the bladder organoid wall are refractory to antibiotic clearance. (**A**) Schematic of the microinjection protocol. Bacteria are injected via a glass microcapillary into the lumen of individual organoids, which are resuspended in a collagen matrix for live-cell confocal imaging. (**B-E**) Snapshots from time-lapse imaging of an infected organoid over 9 hours. (**B**) White arrowhead indicates the organoid boundary. Yellow arrowhead indicates bacteria in the lumen. The bacterial volume within the organoid increases rapidly during the initial growth phase (**B, C**, 0-165 min from the start of the experiment) and occurs predominantly within the lumen. At ca. 165 min, a 10X-MIC dose of ampicillin was added to the extracellular growth media. (**D**) Intra-organoid bacterial volume, quantified by bacterial fluorescence, decreases due to bacterial killing during this period (ca. 165-345 min). Subsequently, the antibiotic was withdrawn (recovery phase ca. 345-525 min) and bacterial regrowth appears to be restricted to the organoid wall (**E**, see supplementary movie SMov1). In contrast, panels (**F-I**) highlight an isolated example where bacterial volume initially decreases (compare **H** vs **G**) but growth subsequently resumes (**I**) even in the presence of the antibiotic (see supplementary movie SMov2). (**J**) Time profiles for intra-organoid bacterial volume over the entire course of the experiment for the organoids in panels **B-E** (squares) and **F-I** (circles), respectively. Shaded region represents the period of antibiotic treatment. (**K-M**) Characterization of bacterial growth dynamics at different stages of infection. (**K, L**) Representative plots of bacterial volume vs. time reveal that growth is exponential during the pre-ampicillin phase (**K**, n = 11 organoids) and decay is exponential during ampicillin treatment (**L**, n = 10 organoids). In some cases, bacterial volume declines immediately after ampicillin administration (**L**, brown lines), while in others, growth continues for some time before declining (**L**, magenta lines). (**M**) Scatter plot for the growth rate of the bacterial volume within the bladder organoids before (n = 124 organoids) and after (n = 58 organoids) ampicillin exposure. The growth rate before ampicillin treatment is significantly faster than the regrowth rate after ampicillin treatment (p<1E-15, Mann-Whitney Test). Red lines represent the median values. Scale bars, 50 μm in (**B-I**).

Addition of ampicillin (Fig. S3B) to the medium surrounding the organoids rapidly reduces the bacterial volume within the organoid (Fig. 2D, H), consistent with the bactericidal nature of the antibiotic. In most cases, bacterial growth resumed only after ampicillin removal (Fig. 2E). Unexpectedly, in two out of the 124 organoids studied, slow bacterial growth continued even in the presence of the antibiotic (Fig. 2I). Representative time profiles for the growth kinetics of UPEC within the organoids for the period before, during, and after ampicillin administration are shown in Fig. 2J, S3C, which capture these two distinct bacterial growth kinetics in the presence of ampicillin.

A plot of bacterial volume over time within infected organoids confirms that growth is exponential (Fig. 2K) with a median growth rate of 0.017 min^-1^ (Fig. 2M), corresponding to a doubling time of 41.5 minutes, in agreement with measurements of growth in the mouse bladder(Justice et al., 2004; Scott et al., 2015). During ampicillin treatment, the bacterial volume initially increases before plateauing due to filamentation of the bacteria, consistent with the mode of action of ampicillin (Fig. 2L). Both the fluorescence intensity and the intra-organoid bacterial volume subsequently decline, with a median killing rate of 0.019 min^-1^ (Fig. S3D). Compared to growth before ampicillin treatment, regrowth after ampicillin washout is significantly slower, with a median growth rate of 0.003 min^-1^, corresponding to a doubling time of 226.8 minutes (Fig. 2M). To rule out the possibility that delayed killing is due to insufficient or delayed drug penetration, we measured the diffusion kinetics of a 0.2 μM solution of lucifer yellow (a fluorescent hydrophilic cell-impermeable dye whose molecular weight is comparable to ampicillin) administered in the medium surrounding the organoids (Fig. S3E, F). Mean fluorescent intensity values in the intraorganoid volume reached a maximum value of ca. 30% of the external concentration within 15-30 minutes of the addition of the dye (Fig. S3F). These results demonstrate that bacteria within the organoids are rapidly exposed to high concentrations of antibiotic in our experimental protocol.

Interestingly, the spatial distribution of bacterial regrowth after ampicillin washout is very different from the lumenal growth observed after microinjection (cf. Fig. 2E vs. 2B, C and Fig. 2I vs. 2F, G), being localized to the bladder wall. At later time points (Fig. 2E), the magenta staining in the interior of the organoid is an artefact of the segmentation pipeline (Fig. S4) and is likely due to expansion of the lumen due to the accumulation of cell debris. We confirmed that regrowth originated from individual bacteria within the organoid wall that survived antibiotic treatment at both 10X MIC (Fig. S5A, B, see supplementary movie SMov4) and 2X MIC (Fig. S5C, D, see supplementary movie SMov5). These results suggest that, whereas the lumen or bladder volume may be the preferred site of growth prior to antibiotic exposure, the bladder wall offers a more protective niche for bacterial survival and regrowth after antibiotic treatment.

### Neutrophils swarm towards intra-organoid UPEC with three distinct migratory profiles

Organoids are powerful systems to study immune cell responses *in situ* (Nikolaev et al., 2020; Sachs et al., 2019; Yuki et al., 2020), and, in conjunction with long-term live-cell imaging, can be used to visualize the spatiotemporal dynamics of immune cell responses. Peripheral innate immune cells such as neutrophils have been shown to be the first responders in the early phases of bladder infection in the mouse model (Haraoka et al., 1999). We therefore added Ly6G+ (Fig. S6A-C), CD11b+ (Fig. S6D-F) murine bone marrow-derived neutrophils to the collagen matrix surrounding bladder organoids immediately after microinjection of the organoid lumen with UPEC. Neutrophils were pre-labeled with CellTracker™ dye to enable identification (Fig. S6B, E) and incubated with murine granulocyte colony stimulating factor (G-CSF) during the co-culture to promote maturation. Live-cell imaging revealed three distinct patterns of neutrophil migration dynamics (see supplementary movie SMov6, SMov7, SMov8). To quantify the dynamics of migration, we used an image analysis pipeline described in greater detail in Fig. S7. Briefly, the organoid surface was identified as described earlier in Fig. S4 and then used as a mask to identify the intraorganoid bacteria, intraorganoid neutrophils, and extraorganoid neutrophils. These volumes were then segmented into surfaces whose properties were measured using standard algorithms in Imaris.

In most cases, neutrophils surrounding an infected organoid migrate towards and accumulate in the lumen, forming aggregates or swarms (Fig. 3A1-3A3, 3B1-3B3). Neutrophil migration into the organoid lumen is consistently accompanied by a sharp reduction in bacterial volume (Fig. 3A2-3A5, Fig. 3B2-3B5, Fig. 3C2-3C5). In some cases, the neutrophil swarms remain within the lumen for >30 minutes (Fig. 3A3-3A5), characteristic of *persistent swarms* (Kienle and Lämmermann, 2016; Lämmermann et al., 2013; Reátegui et al., 2017). In other cases, the swarm rapidly disaggregates after clearance of bacteria within the organoid (Fig. 3B3-3B5), characteristic of *transient swarms*. In a third category, large intra-organoid aggregates of neutrophils do not form; rather, neutrophil numbers within the organoid fluctuate in response to bacterial numbers. We classified this third category, with fluctuating neutrophil numbers inside the infected organoid, as *dynamic swarms* (Hopke et al., 2020). Additional examples for each type of migratory behavior are shown in Fig S9A-C (see supplementary movies SMov9, SMov10, SMov11).

**Figure 3.**
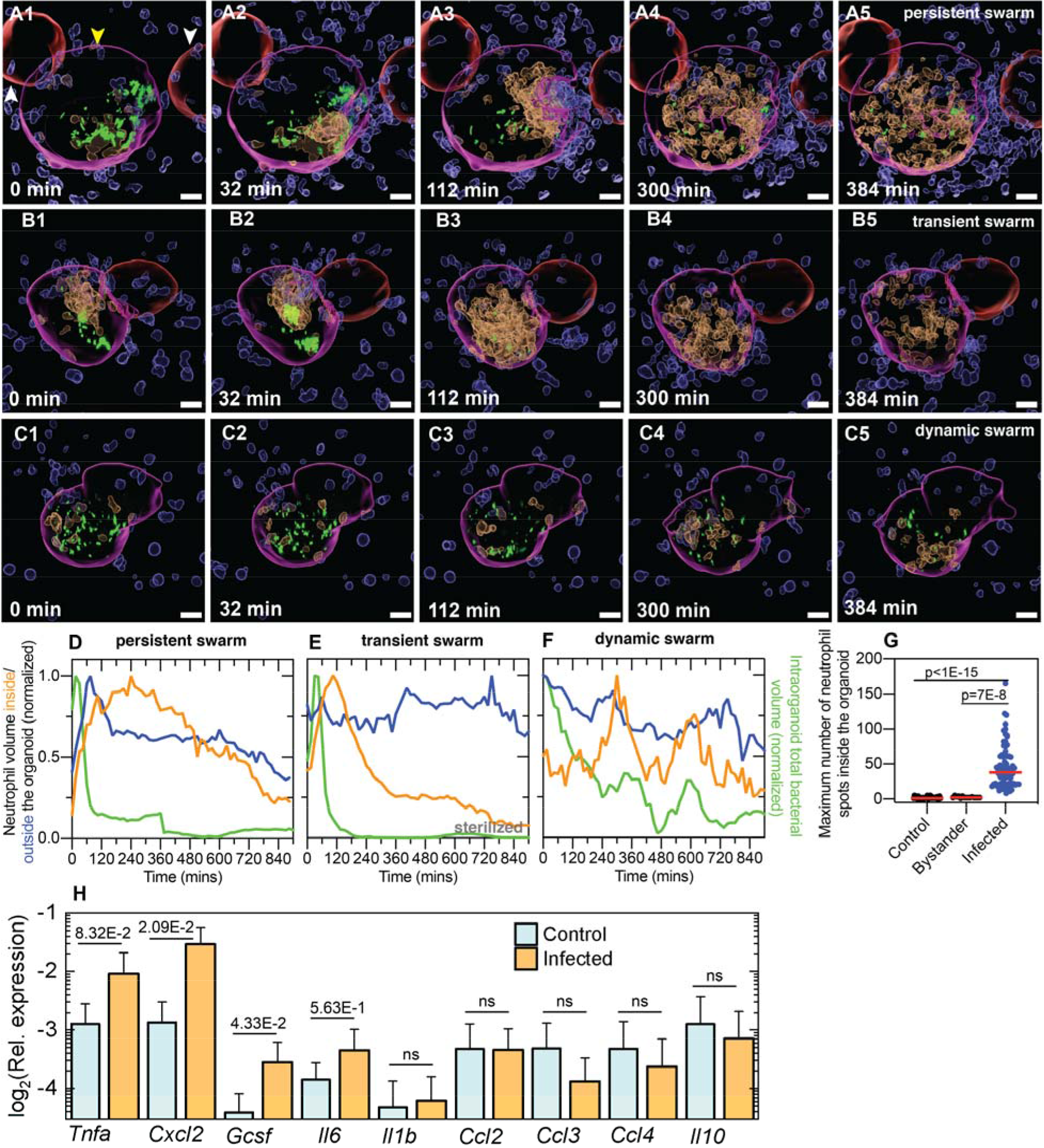
Neutrophil swarming dynamics in response to bacterial infection. Characterization of neutrophil migration dynamics by time-lapse imaging of infected mouse bladder organoids. In all panels, surfaces of intra-organoid (amber) and extra-organoid (blue) neutrophils and infected (magenta) and uninfected (red) organoids were generated using a Bitplane Imaris analysis pipeline. UPEC (green) are shown without processing to identify individual bacteria. (**A1-A5**) **Persistent swarm**. Snapshots show the migration of neutrophils into the lumen of an infected organoid (yellow arrowhead) but not the adjacent uninfected organoid (white arrowhead). A neutrophil swarm forms around the bacteria in the lumen and persists over the course of the experiment. Solitary bacteria in the bladder wall (**A5**) appear to be refractory to clearance. Images are presented in a perspective view. (**B1-B5**) **Transient swarm**. The image series shows two consecutive cycles of neutrophil swarm formation and disaggregation in response to intra-organoid bacterial growth. (**C1-C5**) **Dynamic swarm**. The image series shows rapidly fluctuating neutrophil numbers within the organoid without the formation of large aggregates. (**D-F**) Time profiles of the relative volumes of neutrophils within the organoid, neutrophils surrounding the organoid, and bacteria within the organoid for the three profiles presented in (**A**-**C**), respectively. In each case, the volume is normalized to the maximum volume during the course of the experiment. Persistent and transient swarms are characterized on the basis of neutrophil reverse migration rates. (**G**) Scatter plots of the maximum number of intra-organoid neutrophils during the experiment in uninfected control organoids, uninfected bystander organoids, and infected organoids provides clear evidence of directed migration into infected organoids. P<1E-15 comparing control organoids (n = 30) and infected organoids (n = 75). P=7E-8 comparing bystander organoids (n = 15) and infected organoids (n = 75). (**H**) Plot of the expression of inflammatory cytokines and chemokines relative to *Gapdh* in uninfected and infected organoid populations. The bar represents the mean value, and the error bars represent the standard deviation from n = 4 biological replicates for each condition; each replicate comprises 75-100 infected organoids within a larger population of 750-1,000 organoids. P-values were calculated using Kruskal-Wallis ANOVA Test. Red lines represent median values. Scale bars, 20 μm in (**A1-C5**).

Additional verification of the inward migration of neutrophils is afforded by an examination of the shapes of these cells, as neutrophils surrounding the organoid are predominantly spherical in shape, whereas migratory neutrophils adopt elongated shapes (Fig. S10C). Importantly, neutrophils do not migrate towards uninfected “bystander” organoids, as shown in Fig. 3A, 3B (colored red) and quantified in Fig. S10A, S10B, which confirms that neutrophils can discriminate between uninfected and infected organoids and direct their movement specifically towards infected organoids.

These neutrophil dynamics are better understood from time profiles of the neutrophil volume within the organoid (amber), the neutrophil volume surrounding the organoid (blue), and the intra-organoid bacterial volume (green), each normalized by the maximum value attained over the time period of the experiment. Data for each parameter are shown in Fig. 3D-F corresponding to Fig. 3A1-A5, 3B1-B5, and 3C1-C5, and Fig. S9D-F corresponding to Fig. S9A-C, respectively; absolute numbers are shown in Fig. S8 and Fig. S9G-I. Infected bladder organoids therefore successfully reproduce the range of neutrophil migratory responses reported from intravital imaging studies, and bacteria within the lumen are effectively cleared by incoming neutrophils independent of their migratory profile. We characterized mixed populations of infected and bystander organoids relative to uninfected organoids at ca. 4 hours post infection by measuring the expression of genes that could stimulate leukocyte migration and genes that are known to be upregulated during UPEC infection *in vivo* (Fig. 3H). We found that, expression of the inflammatory cytokine *Tnfa* (Yu et al., 2019) the neutrophil maturation factor *Gcsf* (Ingersoll et al., 2008; Semerad et al., 2002), and the neutrophil chemoattract *Cxcl2*, (Sundac et al., 2016) were more highly expressed in infected/bystander organoids compared to uninfected controls (Sundac et al., 2016). A small but statistically insignificant increase was observed in the expression of the inflammatory cytokine *Il6*, and no changes in expression were observed for macrophage-specific chemokines such as *Ccl2* or in the expression of *Ccl3* and *Ccl4*, which also stimulate neutrophil migration but are typically produced by resident macrophages (Mariano et al., 2020; Schiwon et al., 2014). These observations are consistent with the cellular composition of the organoids as well as reports from the mouse model, where UPEC infection does not increase *Ccl3* expression and leads to increased *Ccl4* expression only at later stages of infection (Ingersoll et al., 2008). These infection-driven changes in host-cell gene expression could explain, in part, the directed migration of neutrophils towards infected organoids.

### Bacteria within the wall of the bladder organoid are refractory to clearance by neutrophils

The rapid clearance of lumenal bacteria by migratory neutrophils provides an opportunity to observe niches where bacteria may be protected from neutrophil attacks, such as intracellular bacterial communities (IBCs) (Justice et al., 2004). We observed IBC-like structures in a subset of infected organoids, reinforcing the validity of the organoid model. Hereafter, for simplicity, we shall refer to these structures as “IBCs” although we acknowledge that the IBC-like structures observed in the organoid model may differ in some respects from the *bona fide* IBCs that form in the bladder wall of infected animals. Fig 4A1-3 shows an example of an organoid IBC that forms and persists despite the presence of a persistent swarm of neutrophils within the organoid that successfully clears the bacteria from the organoid lumen. Snapshots from the raw images at the corresponding time points are shown in Fig. S11 to confirm that the IBC is formed in a uroepithelial cell. An additional example in Fig. S12 confirms that these structures are formed in CK8+ cells. Live-cell imaging revealed the IBC to be a dynamically fluctuating structure; for example, in the 15-minute interval between Fig. 4A3 and Fig. 4A4 (see supplementary movie SMov12), the IBC begins to shed bacteria, which are rapidly taken up by nearby neutrophils. The vast majority of the bacteria released from the IBC are processed and killed within 30 minutes, as evidenced by loss of YFP fluorescence (Fig. 4A5). In contrast, Fig. 4B1-5 (see supplementary movie SMov13) shows another example of an IBC that forms in the presence of a persistent neutrophil swarm (white arrowhead, Fig. 4B1-2), but in this case the bacteria shed from the IBC are spread by the neutrophils to different regions of the bladder epithelium (Fig. 4B3-4, cyan arrowheads). Some of these bacteria persist as isolated bacteria within the organoid wall and appear to resist clearance by neutrophils (cyan arrowheads, Fig. 4B5).

**Figure 4.**
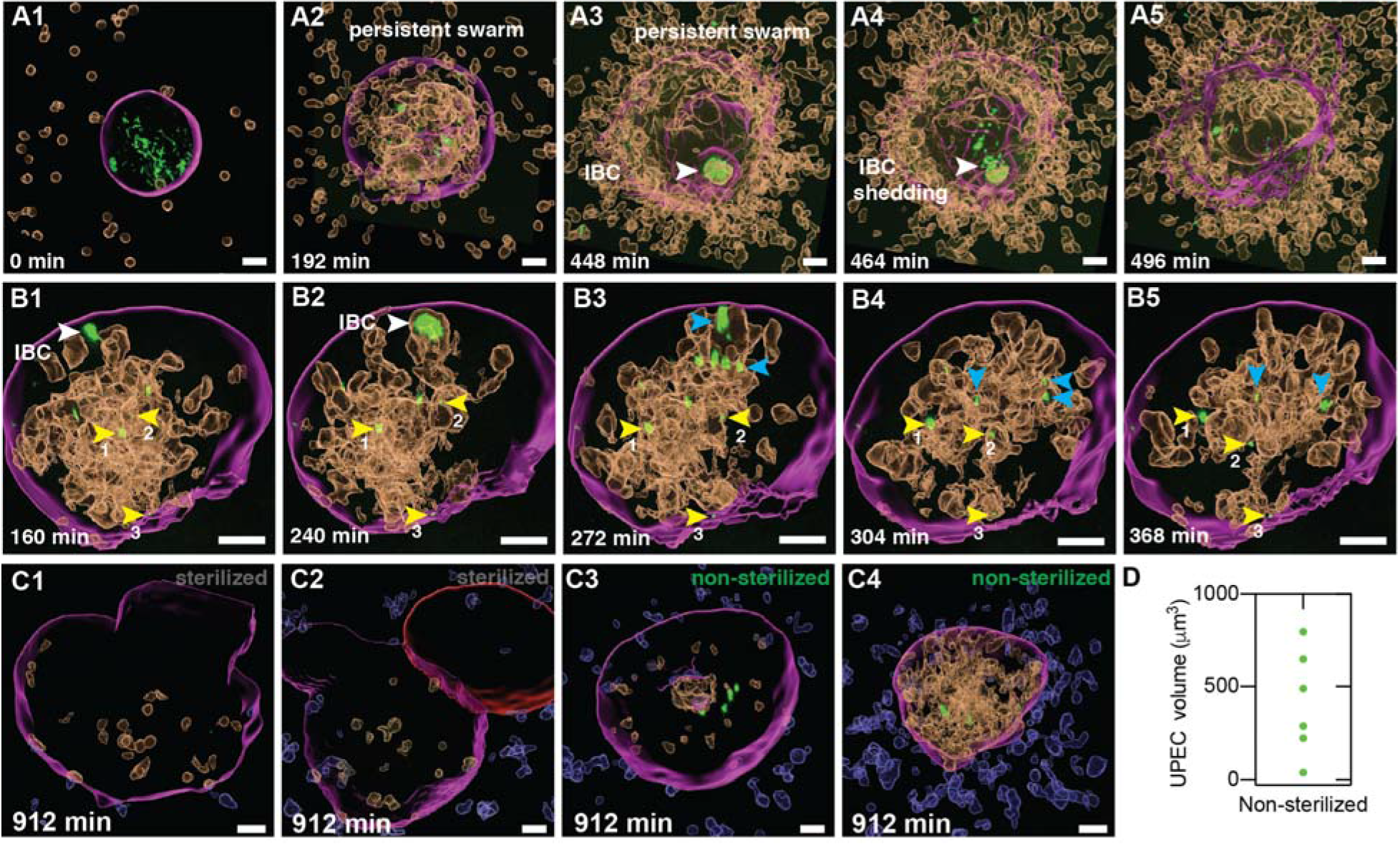
Bacteria within the bladder organoid wall are protected from clearance by neutrophil swarms. (**A1-A5, B1-B5**) Two examples of IBCs that are protected from neutrophil-mediated clearance. Shedding of bacteria from an IBC (**A4, A5** and **B2, B3**) is followed by two outcomes: phagocytic uptake and rapid clearance (**A5**), or dispersal to other niches in the bladder epithelium (**B5**, cyan arrowheads). Three examples of isolated solitary bacteria in the bladder wall (labelled 1, 2 3) that are refractory to clearance throughout the time series in (**B1-B5**) are indicated by yellow arrowheads and tracked within the organoid over this time period. (**C1-C4**) Snapshots at ca. 15 hours after infection and addition of neutrophils. (**C1, C2**) Examples of bacterial clearance from organoids following neutrophil treatment. (**C3, C4**) Examples of bacterial persistence in organoids where the neutrophil swarm disperses (**C3**) or persists up to 15 hours post-infection (**C4**). (**D**) Quantification of the total bacterial volume at 15 hours post-infection in case of non-sterilized organoids (n = 6). Scale bars, 20 μm in (**A1-A5**), (**B1-B5**), and (**C1-C4**).

Unexpectedly, at early time points we also identified spatially isolated subpopulations of bacteria, comprising individual cells or small clusters (the threshold of detection was set at 10 μm^3^), located within the organoid wall (yellow arrowheads, Fig. 4B1). Hereafter, we refer to these subpopulations as “solitary” bacteria to distinguish them from “communal” bacteria within IBCs. The high spatiotemporal resolution afforded by confocal imaging allowed us to track three such examples, labelled ‘1’ through ‘3’, over the entire course of the time series shown in Fig. 4B1-5. Our results confirm that solitary bacteria are located within the organoid wall throughout the time course, and they appear to be refractory to neutrophil-mediated clearance (Fig. 4B5, yellow arrowheads and corresponding labels). The number of solitary bacteria was enhanced by the addition of some of the bacteria shed by the ruptured IBC (cyan arrowheads in Fig. 4B5). Even 15 hours after addition of neutrophils, solitary bacteria persist within some organoids (Fig. 4C3, 4C4, 3C and analysis in Fig. 3D, 3F, S8C, S9D, S9G), whereas other organoids are successfully sterilized by the neutrophils (Fig. 4C1, 4C2, and analysis in Fig. 3E, Fig. S8B, and Fig. S9E-I). These two outcomes (persistence and sterilization) occur with roughly equal frequency (Fig. 4D). Live-cell imaging confirmed that bacteria shed from IBCs contribute only a small portion of these persistent bacteria.

These results demonstrate that niches within the deeper layers of the bladder epithelium, below the superficial layer of umbrella-like cells containing IBCs, can also harbour subpopulations of bacteria that are resistant to clearance by antibiotics or neutrophils. In the latter case, bacterial persistence is independent of the migratory profile of the neutrophil swarm (Fig. 4C3, 4C4). We attempted to localize these persistent bacteria with greater precision, but the spatial resolution of confocal microscopy proved to be insufficient to identify unambiguously the boundaries of the bladder lumen and the precise location of individual bacteria within the organoid wall. We therefore turned to electron microscopy, which provides much higher spatial resolution than optical microscopy.

### Volumetric electron microscopy reveals five distinct bacterial niches within infected organoids

Technical developments in scanning electron microscopy now permit volumetric imaging of large samples using serial block face scanning electron microscopy (SBEM) (Denk and Horstmann, 2004; Hoffman et al., 2020; Maclachlan et al., 2018). High-contrast staining used for the preparation of SBEM samples densely labels cell membranes, which is useful for localizing bacteria within tissue. We used this imaging method to capture an entire infected organoid at ca. 6 hours after microinjection of UPEC and addition of neutrophils. The organoid was imaged with optical microscopy (Fig. S13A) prior to staining and resin embedding. It was then mounted inside the scanning electron microscope and serial images of the entire structure were collected at a lateral resolution of 30 nm with 100 nm sections separating each image; examples of such slices are shown in Fig. S13B1-B4. The imaging parameters allowed us to identify cell membranes and to localize all bacteria within the organoid. In addition, optical microscopy imaging prior to embedding provided a 3D map of the organoid, which facilitated the identification of fluorescently labelled cells such as neutrophils in the final EM image series. We identified a total of 2,938 bacteria and classified them into five distinct subpopulations according to their locations within the organoid (Fig. 5A-E, Fig. S13B1-B4, see supplementary movie SMov14).

**Figure 5:**
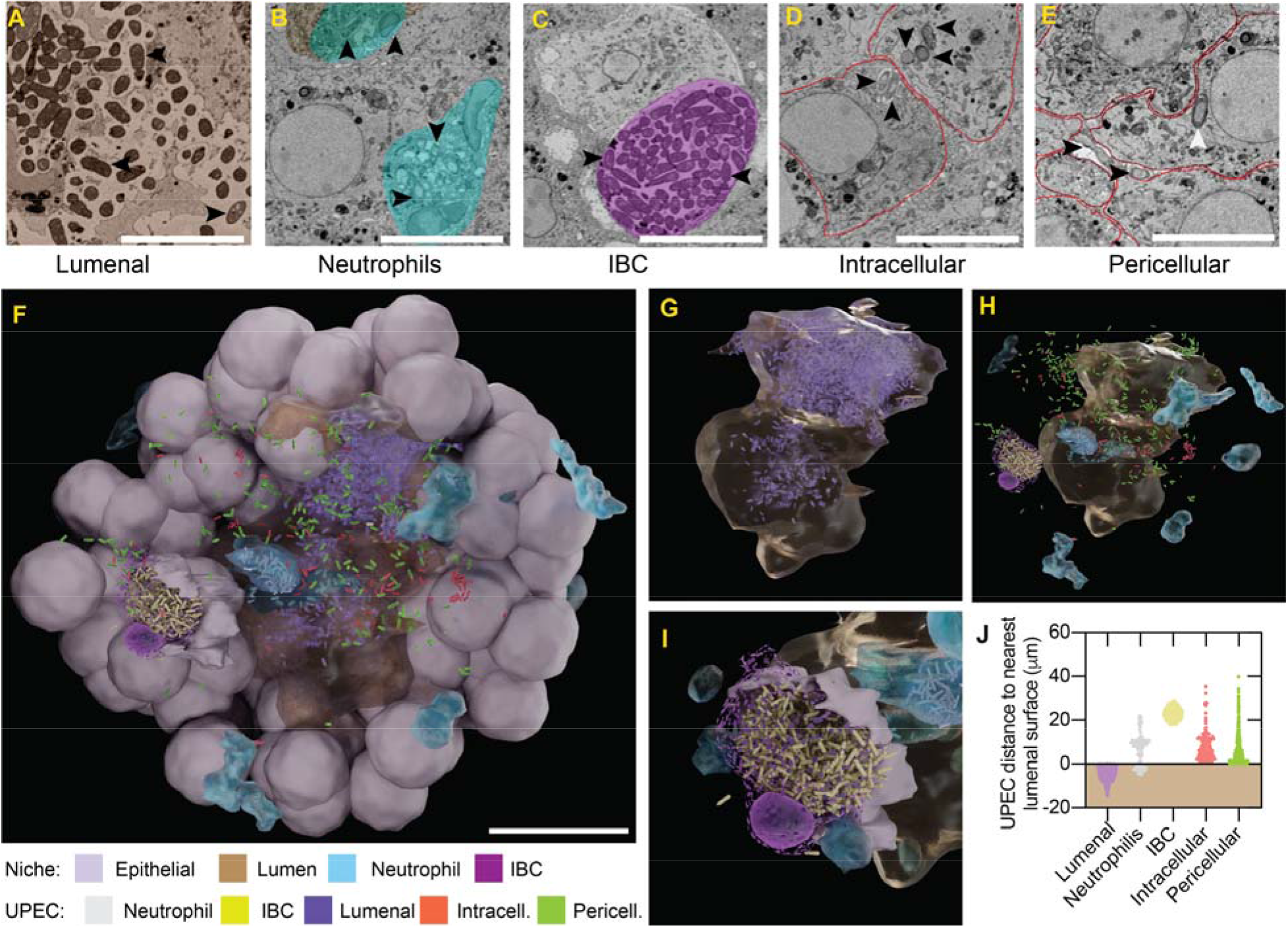
Volumetric electron microscopy reveals five distinct bacterial niches within an infected organoid. (**A-E**) Serial-block face-scanning electron microscopy (SBEM) snapshots of an infected organoid at ca. 6 hours after microinjection of UPEC and addition of neutrophils. Five distinct niches were identified. Bacteria within each niche are indicated by black arrowheads. (**A**) Extracellular bacteria within the organoid lumen (brown; n = 1,821). (**B**) Bacteria within neutrophils (light cyan) that can be either in the lumen or in the bladder wall (n = 141). (**C**) Bacteria within an IBC (n = 398). The area of the IBC is shown in purple. (**D**) Individual intracellular bacteria within the cytoplasm of epithelial cells (cell boundaries in red; n = 111). (**E**) Individual pericellular bacteria located between epithelial cells (cell boundaries in red; n = 467). An additional example of an intracellular bacterium is indicated with a white arrowhead. (**F**) Model derived from the serial electron microscopy images of the entire organoid in which all bacteria and cells were plotted. Image shows the interior of the organoid, revealing the lumen (brown), as well as epithelial cells (grey) and neutrophils (cyan). Bacteria located within the five niches corresponding to (**A**-**E**) are colored violet (lumenal), grey (neutrophil), yellow (IBC), red (intracellular), and green (pericellular). The width of the organoid is 85 μm. (**G**) Zoom shows the extracellular bacteria in the lumen (brown). (**H**) Zoom shows all other bacterial classifications around the lumen. (**I**) Zoom shows the bacteria within the IBC, which appear to be protected from clearance by the surrounding neutrophils. (**J**) Scatter plot of the distance of bacteria within each subpopulation from the lumenal surface. Lumenal bacteria are located entirely within the lumen (shaded brown region). Scale bars, 5 μm in (**A-E**) and 20 μm in **F**.

The majority (62%, n = 1,821) of the bacteria are extracellular and located within the organoid lumen (Fig. 5A, Fig. S13B1). Smaller bacterial fractions are located either as solitary bacteria within the cytoplasm of bladder epithelial cells (3.8%, n = 111) or within the IBC (13.5%, n = 398) (Fig. 5C, D, Fig. S13B2-B3). The small fraction of bacteria located within neutrophils (4.8%, n = 141) is probably an underestimate, since bacteria are typically degraded after phagocytic uptake (Fig. 5B, Fig. S13B2). Bacteria within the neutrophils have an altered morphology, likely due to exposure to antimicrobial stresses within the neutrophil. Unexpectedly, we identified a fifth subpopulation of solitary bacteria, which we term “pericellular” (15.9%, n = 467), that is located in between bladder epithelial cells within the organoid wall (Fig. 5E, Fig. S13B4). To the best of our knowledge, this pericellular subpopulation of solitary bacteria has not been reported previously, presumably due to the difficulty of whole-bladder imaging with sufficient spatial resolution to identify individual bacteria located between host-cell membranes.

A view inside a model of the organoid, created from a 3D map of coordinates of all cells and bacteria, shows the arrangement of the different bacterial subclasses inside (Fig. 5F). The bacterial population within the irregularly shaped lumen clusters towards the first quadrant (zoom in Fig. 5G), likely indicating the site of the injection. Intracellular and pericellular solitary bacteria can be found scattered throughout all areas of the organoid wall (zoom in Fig. 5H). Individual neutrophils can be observed in all areas of the organoid wall as well as the lumen (Fig. 5H), and some of these neutrophils contain bacteria (Fig. S13F). Bacteria within the IBC form a tight cluster below the nucleus of the cell hosting the IBC (Fig. 5F). The protection against phagocytic uptake conferred by intracellular localization is evident from the fact that the IBC remains intact despite the infected cell being surrounded by multiple neutrophils. A plot of the shortest distance of each subpopulation to the organoid lumen is shown in Fig. 5J. It is noteworthy that intracellular and pericellular solitary bacteria are found within the deeper layers of the organoid that express intermediate and basal cell markers (Fig. 1C).

We asked whether the intracellular and pericellular solitary subpopulations are phenotypically different from bacteria in the organoid lumen. UPEC has been shown to markedly alter flagellar expression and cell shape during bladder infection (Anderson et al., 2004; Lane et al., 2007; Wright et al., 2007). We therefore used immunofluorescence to probe flagellin expression *in situ*. Because paraformaldehyde fixation tends to degrade the fluorescence signal from bacterial YFP, we used immunostaining against lipopolysaccharide (LPS) and a mask of the organoid shape to label intra-organoid bacteria (cyan) and bacteria surrounding the organoid (green) in Fig. 6A. The corresponding image with anti-flagellin immunostaining (Fig. 6B) shows that a majority of intra-organoid bacteria have low or no detectable flagellin expression. The specificity of the anti-flagellin antibody was confirmed using a flagellin-deficient Δ*fliC* strain (Fig. S14A). We applied a threshold on the basis of volume to the intra-organoid bacteria clumps and found that large bacterial clumps (volume greater than 1,000 μm^3^) express very low flagellin levels (Fig. 6D), whereas smaller clusters and individual bacteria retain intermediate to high levels of flagellin expression (Fig. 6C, Fig. S14B). Thus, bacteria that continue to express flagellin are predominantly single cells or small clusters located within the organoid wall; both observations suggest that these subpopulations correspond to the intracellular or pericellular subpopulations of solitary bacteria identified in our serial electron micrographs. This point is strengthened by our observation that bacteria within the IBC have severely reduced flagellar expression, as shown previously in infected mice (Wright et al., 2005), whereas single bacteria surrounding the IBC retain high-level flagellar expression (Fig. 6E).

**Figure 6:**
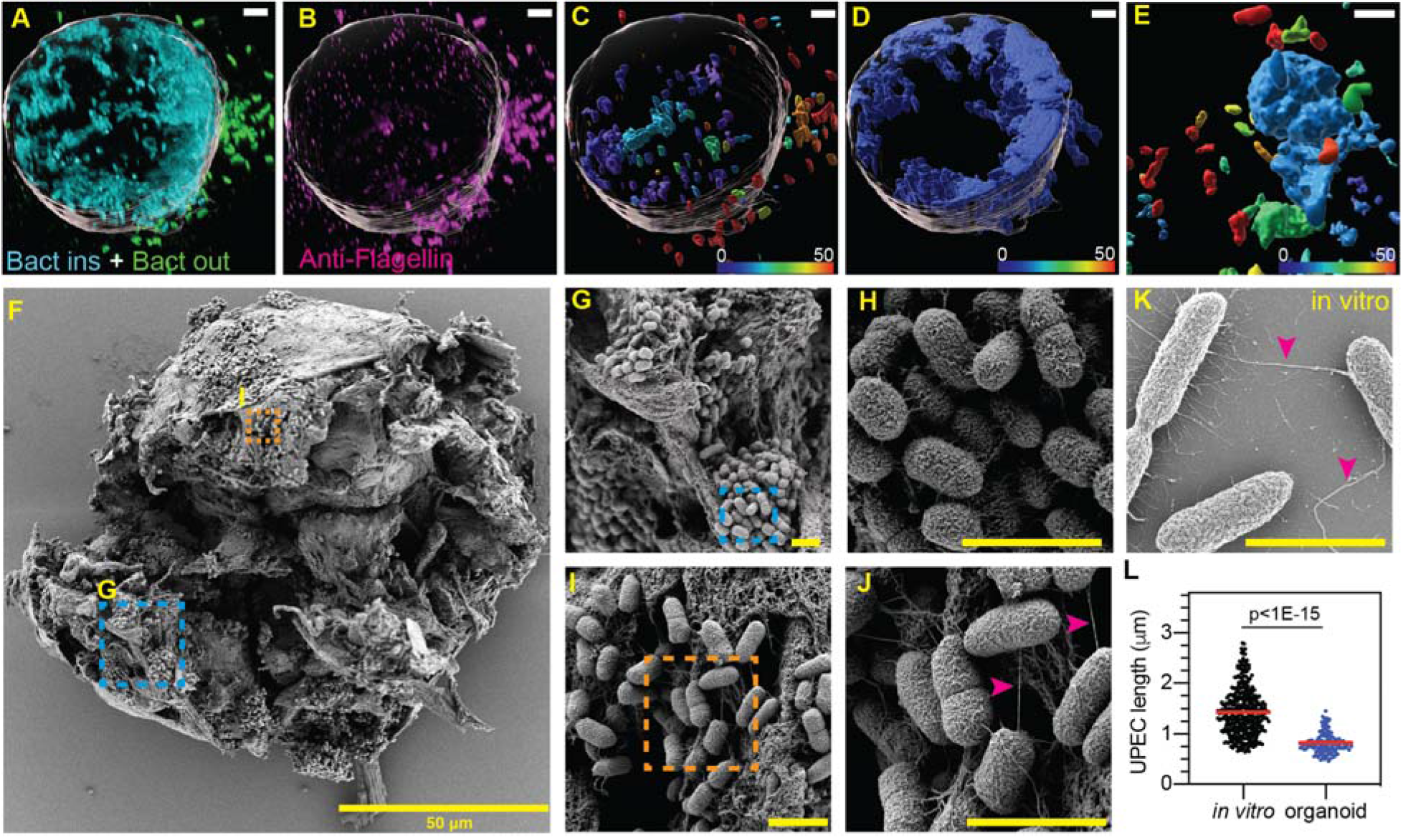
Solitary bacteria within the bladder organoid wall are rod-shaped and flagellated whereas bacteria within dense clusters are coccoid and non-flagellated. (**A-D**) Perspective views of an infected organoid (surface indicated in grey) at ca. 6 hours after microinjection with UPEC. (**A**) Bacteria inside (cyan) and outside (green) the organoid were identified by immunostaining for bacterial LPS. (**B**) Flagellated bacteria (magenta) were identified by immunostaining for bacterial flagellin. Analysis of co-expression of LPS and flagellin for individual bacteria (**C**) or large clumps of bacteria, which are predominantly restricted to the lumen (**D**). Bacteria are color-coded according to the intensity of the flagellin signal. Flagellated bacteria are predominantly observed outside the organoid (**C**) but can also occur as individual solitary cells within the organoid wall. (**E**) Bacteria within an IBC have low-level flagellin expression, whereas individual bacteria surrounding the IBC retain high-level flagellin expression. (**F**) Scanning Electron Microscopy (SEM) image of an infected organoid at 6 hours after microinjection with UPEC. Ruptured organoids reveal multiple areas of bacterial growth. Zooms for two representative areas are shown in (**G-J**). Bacteria in (**I**) are loosely packed, rod-shaped, and express flagella (**J**, magenta arrowheads). Bacteria in (**G**) are tightly packed, coccoid-shaped, and do not express flagella (**H**). (**K**) SEM image of UPEC grown in axenic culture. Cells are rod-shaped and flagellated (magenta arrowheads). (**L**) Size distributions for bacteria grown to stationary phase in axenic culture and bacteria within tightly packed clusters in the organoid. Both populations were imaged by SEM. Bacteria in axenic cultures (n = 370) are longer than intra-organoid bacteria (n = 136) (p<1E-15, calculated using Mann-Whitney Test). Scale bars, 5 μm in (**A-E)** and 2 μm in (**G-K)**.

Scanning electron microscopy (SEM) is a powerful technique to examine surface morphology and overall bacterial shape. We ruptured open an infected organoid using a tungsten needle (Heo et al., 2018) at ca. 6 hours after microinjection with UPEC to gain access to the interior of the organoid (Fig. 6F). We found that the majority of bacteria accessible to SEM imaging are located within tight clusters, suggestive of IBC growth (Fig. 6G, H). These bacteria are coccoid in shape and are not flagellated (Fig. 6H), consistent with previous findings in infected mice (Anderson et al., 2003; Justice et al., 2004). In contrast, we identified a small subpopulation of bacteria in a different spatial location within the organoid that retain flagellar expression (Fig. 6I, J). These observations are consistent with data in Fig. 6C, D. We verified that bacteria from a stationary-phase axenic culture retain flagellin expression (Fig. 6K) and are longer than intra-organoid bacteria (Fig. 6L). The bladder organoid model is therefore able to recapitulate characteristic morphological features of bacteria within IBCs. It also reveals that the bladder wall comprises distinct niches, including some in which the bacteria are rod-shaped and retain flagellin expression. These observations highlight the utility of organoids as a tool to obtain a more comprehensive picture of UPEC phenotypic variants that arise during the course of infection.

## Discussion

The mouse model of UPEC infection faithfully reproduces key features of UPEC pathogenesis in humans (reviewed in Flores-Mireles *et al*. (Flores-Mireles et al., 2015) and Hung *et al*. (Hung et al., 2009)), but the complexity of this model poses difficulties for live-cell imaging with high spatiotemporal resolution. There is also a paucity of tractable *in vitro* models that reproduce the complex 3D stratified architecture of the bladder wall and key features of the UPEC pathogenesis cycle, such as bacterial persistence and regrowth after antibiotic treatment. In one early study, Smith *et al*. (Smith et al., 2006) reported that a human cancer cell line could be induced to establish stratified uroepithelial layers on collagen beads by exposure to shear stress. However, this model lacks a lumen where bacteria can be introduced in a topologically correct manner. Andersen *et al*. (Andersen et al., 2012) developed a model in which a virus-immortalized human bladder cell line is cultured under flow to mimic micturition, but this model does not include a stratified uroepithelium or the ability to introduce additional cell components, such as immune cells. More recently, Horsley *et al*. (Horsley et al., 2018) succeeded in growing primary human uroepithelial cells as stratified layers on Transwell® inserts, although in this model stratification is non-uniform and it is difficult to reproduce immune cell dynamics or track them using live-cell imaging. Although other stratified bladder epithelium models have been reported (Cattan et al., 2011; Chabaud et al., 2017; Halstead et al., 2017; Suzuki et al., 2019), to the best of our knowledge, none of these models has been adapted for infection studies.

Here, we show that bladder organoids embedded in a collagen matrix fulfill all of these requirements: the ability to track bacterial growth within the organoid wall and within a central lumen that mimics the bladder volume; the ability to add and remove soluble compounds, which we exploit here for real-time analysis of bacterial responses to antibiotic treatment; and the ability to study the directed migration of immune cells into organoids in response to infection. Organoids are amenable to long-term live-cell imaging and offer a medium-throughput format with the flexibility to combine different cell types, such as uroepithelial cells and immune cells derived from different strains of genetically modified mice. These combinations are not possible in simple monolayer systems (Andersen et al., 2012) or in previously reported complex stratified systems (Horsley et al., 2018; Smith et al., 2006). In these respects, the organoid model also offers advantages compared to experiments with explanted bladder tissue (Justice et al., 2004, 2006), Shortcomings of the organoid model in its current form include the absence of resident immune cells (Mariano et al., 2020; Schiwon et al., 2014) or vasculature (Homan et al., 2019; Mansour et al., 2018), the accumulation of cell debris in the lumen, and the inability to expose the cells of the umbrella-like layer facing the lumen to urine. Some of these limitations may be addressed with a bladder-chip platform (Sharma et al., 2021), or through the development of appropriate organoid-on-chip systems (Park et al., 2019; Takebe et al., 2017; Zhang et al., 2018).

We leverage these advantages to demonstrate that whereas the bladder organoid lumen is the predominant site of bacterial replication, it is the simultaneous presence of bacteria within the organoid wall that enables infection to resist clearance by antibiotics and neutrophils. Prior to antibiotic treatment, rapid growth of bacteria in the organoid lumen predominates, although widely scattered invasion of bacteria into the bladder wall occurs at the same time. Some of the invading bacteria take up residence within the superficial umbrella-like cells abutting the organoid lumen, where they replicate intracellularly to form IBCs, while others invade into deeper layers of the organoid wall. Bacteria within IBCs are communal, non-flagellated, coccoid-shaped, and intracellular. In contrast, bacteria that invade into deeper layers of the organoid wall are solitary, flagellated, rod-shaped, and may be intracellular or pericellular. Following antibiotic washout, the spatial pattern of bacterial growth is reversed, occurring at scattered sites throughout the organoid wall but only rarely within the lumen. Also, regrowth in the organoid wall after antibiotic treatment is significantly slower than growth in the organoid lumen prior to treatment. In rare cases, we found that bacteria within the organoid wall continue to divide even in the presence of antibiotic. This observation could reflect a niche within the organoid wall where antibiotic penetration is poor, emergence of antibiotic resistance, or the fact that bacterial responses to antibiotics are heterogeneous and dynamic and may, in some cases, involve balanced division and death (Wakamoto et al., 2013). These dynamic and transient phenotypes are difficult to capture in animal models or using conventional static measurements of bacterial numbers such as colony forming units. We conclude that early invasion of solitary bacteria into deeper layers of the bladder wall, concomitant with invasion of superficial umbrella cells and IBC formation, may play an important role in bacterial persistence and relapse following antibiotic treatment.

We observed similar spatially distinct dynamics for bacterial killing by neutrophils. The small volume and spherically symmetric geometry of the organoid model allowed us to image and quantify neutrophil migration dynamics, which would be difficult to achieve in Transwell-based models. We found that UPEC infection of organoids generates a strong and highly directed neutrophil migration response, with three distinct spatiotemporal patterns (persistent, transient, or dynamic) that are reminiscent of descriptions based on intravital imaging of neutrophil responses to bacterial infections (Hopke et al., 2020; Isles et al., 2019; Kienle and Lämmermann, 2016; Lämmermann et al., 2013; Liese et al., 2012; Poplimont et al., 2020; Shannon et al., 2013) relative frequencies of these behaviors may differ *in vivo* due to the presence of a complete resident immune system. We also found that neutrophils migrate out of the lumen of bladder organoids after resolving infection. Regardless of the neutrophil migration pattern, about half of the organoids that we studied were not completely sterilized by neutrophils even after many hours. In the majority of these cases, bacterial survival was restricted to the organoid wall while lumenal bacteria were sterilized.

Although both IBCs in superficial umbrella-like cells and solitary bacteria within deeper layers of the organoid wall appear to be relatively refractory to clearance by neutrophils, we found that the bacteria released when IBCs rupture are rapidly taken up and destroyed by patrolling neutrophils. In a subset of organoids where we did not detect any IBCs, solitary bacteria within deeper layers of the uroepithelium were solely responsible for survival during neutrophil attacks. Although we observed both pericellular and intracellular solitary bacteria by SBEM, it is possible that the intracellular population is more likely than the pericellular population to survive attacks by neutrophils or other resident immune cells. The organoid model may exaggerate the numbers of solitary bacteria due to the lack of resident macrophages or the insufficient maturation and activation of bone marrow-derived neutrophils in the model. Nevertheless, these observations suggest that early and IBC-independent invasion of solitary bacteria into deeper layers of the bladder wall might play an important role in recurrent infections by generating a subpopulation that is refractory to clearance by antibiotics and the host innate immune response.

Solitary bacteria in deeper layers of the bladder wall are phenotypically distinct from bacteria within IBCs, inasmuch as they are rod-shaped and flagellated, whereas bacteria within IBCs are coccoid-shaped and non-flagellated. The latter point is interesting because loss of flagellar expression within IBCs has been reported previously (Wright et al., 2005), whereas flagellar expression has been shown to be important for generating persistent infections in the mouse bladder (Lane et al., 2007; Wright et al., 2005). Our findings suggest that non-flagellated bacteria within IBCs and flagellated solitary bacteria seeded throughout deeper layers of the bladder wall may both contribute to survival during antibiotic treatment and neutrophil attacks. Given the relatively small numbers of solitary bacteria in the deeper layers of the organoid wall, it is unlikely that this subpopulation could be identified using population-averaged measurements, such as transcriptomic analysis of the total bacterial population within the bladder, or by non-exhaustive microscopic imaging of bladders from infected mice.

The small volume of the organoid lumen and the lack of bacterial clearance by micturition might accelerate the formation of IBCs relative to the mouse model. Even so, it is noteworthy that spatially distinct IBCs and solitary bacteria within deeper layers of the organoid wall both appear within hours of infection. This contrasts with the current model of UPEC persistence, which postulates that cycles of formation and rupture of IBCs, resulting in exfoliation of superficial umbrella cells and exposure of underlying layers of the stratified epithelium, are a precursor to invasion of bacteria into deeper layers of the uroepithelium, where they generate “quiescent intracellular reservoirs” (Mysorekar and Hultgren, 2006). At present we cannot rule out the possibility that quiescent intracellular reservoirs and solitary bacteria in deeper layers of the bladder wall, as described here, are phenotypically equivalent. Although these populations seem to arise on different time scales, it may be that the barrier to invasion of the bladder wall is less robust in the organoid model compared to the mouse bladder, permitting earlier colonization of deeper layers. Alternatively, it is possible that early invasion of bacteria into deeper layers of the bladder wall may have been overlooked in previous studies in infected mice due to the difficulty of detecting small numbers of isolated bacteria within a large mass of tissue.

The dynamics of host-pathogen interactions during bladder infections are difficult to capture with high spatiotemporal resolution in conventional animal models. Bladder organoids, being miniaturized and experimentally tractable models of the bladder, are well positioned to generate new insights into UPEC pathogenesis, although it is important to note that organoid models lack the full complexity of the *in vivo* tissue environment. Thus, we view the two models, animal and organoid, as complementary approaches with distinct advantages and disadvantages. Here, we use time-lapse optical microscopy and electron microscopy to demonstrate the existence of solitary subpopulations of intracellular and pericellular bacteria located within deeper layers of the stratified bladder organoid wall, beneath the superficial layer of umbrella cells. These solitary subpopulations appear early in the course of infection, concomitant with the formation of IBCs in the umbrella-cell layer, and they resist elimination by antibiotics and neutrophils. Improved understanding of the physiology of solitary bladder wall-associated bacteria could contribute to the development of new strategies to eliminate persistent bladder infections.

## Acknowledgements

V.V.T gratefully acknowledges support by a Human Frontier Science Program (HFSP) Long-Term Fellowship (LT000231/2016-L) and a European Molecular Biology Organization (EMBO) Long-Term Fellowship (921-2015). This research was supported by grants to J.D.M. from the Swiss National Science Foundation (grant number 310030B_176397) and the National Centre of Competence in Research AntiResist funded by the Swiss National Science Foundation (grant number 51NF40_180541). K.S wishes to thank the entire team of EPFL Bioimaging & Optics Core Facility for their assistance in confocal live cell imaging and post analysis in Bitplane Imaris, Dr. Devanjali Dutta (EPFL) and Stephanie Rosset (at the EPFL Biological Electron Microscopy Facility) for help in optimizing the protocol for rupturing of infected organoids, Dr. Jessica Sordet-Desimoz and Gian-Filippo Mancini (EPFL Histology Core Facility) for assistance with paraffin-embedded slicing of organoids, Dr. Germann Markus (EPFL) for development of the neutrophil isolation protocol, Mikhail Nikolaev (EPFL) for assistance with the collagen gel polymerization protocol, Thomas Simonet (EPFL) for the UPEC Δ*fliC* strain, Alexandre Marcos (EPFL) for optimizing the protocol for microinjection of bladder organoids, and Prof Andrew Oates for kindly providing access to a stereo microscope. The authors thank and credit BioRender.com for the illustrations and schematics used in this manuscript.

## Author Contributions

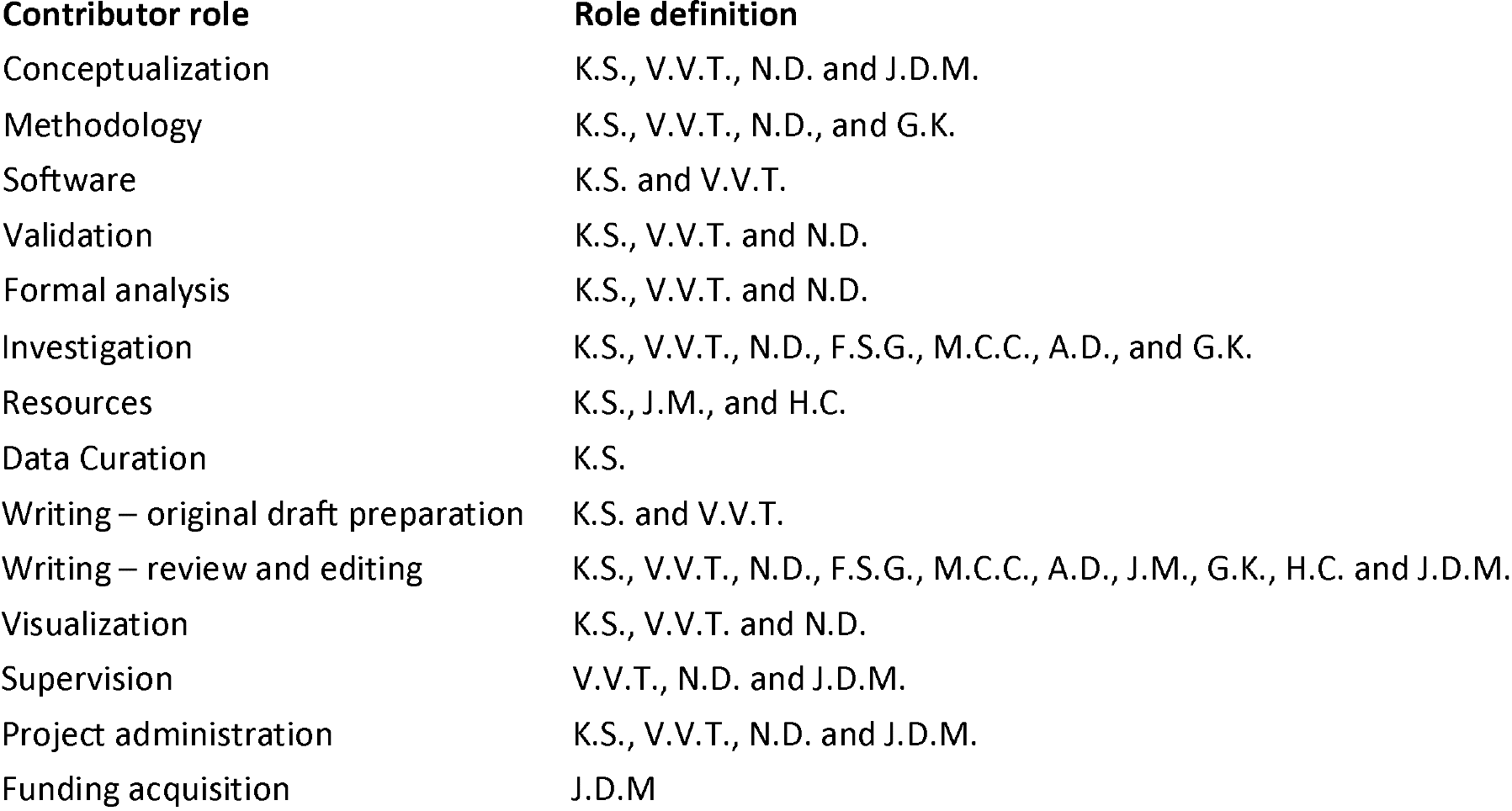

## Materials and Methods

### Generation of mouse bladder organoids from mouse bladder uroepithelial cells

C57BL/6 mice (Charles River Laboratories) or ROSA^MT/MG^ mice (Jackson Laboratories) Mice used for these experiments were housed in a specific pathogen-free facility. All animal protocols were reviewed and approved by EPFL’s Chief Veterinarian, by the Service de la Consommation et des Affaires Vétérinaires of the Canton of Vaud, and by the Swiss Office Vétérinaire Fédéral. Mouse bladder organoids were prepared by selectively isolating lumenal cells from the bladder of following the procedure described in Mullenders *et al*. (Mullenders et al., 2019).

For the generation of a large number of organoids, three female mice at age of four months were were euthanized by CO_2_ overdose. Lumenal uroepithelial cells were isolated by microinjecting approximately 500 μL TryPLE (Gibco) with a 26G needle (Terumo) into the lumen of the bladder. During this procedure, the urinary bladder was clamped at the outlet. Subsequently, the clamped urinary bladder filled with TrypLE solution was immersed in pre-warmed 20 mL Basal Medium (Advanced DMEM/F-12 medium) and placed in a 50 mL falcon tube. The falcon tube containing the bladder was subsequently incubated for 1 hour in a cell culture incubator maintained at 37°C and 5% CO_2_. The lumen of the bladder was then washed twice with basal medium containing 20% heat-inactivated fetal bovine serum (FBS, Gibco) to neutralize the effect of TryPLE. The cell suspension was passed through a 40 μm filter (Fisher) and the cells in the flow-through were centrifuged at 300 g for 5 minutes. These isolated uroepithelial cells were then resuspended in an appropriate volume of Cultrex® Basement Membrane Extract (BME) and seeded as hemispherical domes (40-50 μL in volume) in individual wells of a 24-well plate (NAME). the 24-well plate was placed in a cell culture incubator for 30 minutes and during this period it was maintained in an inverted configuration to promote 3D growth. The solidified hemispherical domes were then surrounded by mouse bladder medium (MBM, (Mullenders et al., 2019)) supplemented with 1X antib acterial/anti fungal solution (Gibco). MBM medium consists of Advanced DMEM/F-12 medium (Thermofisher), 100 ng/mL of FGF10 (Peprotech), 25 ng/mL of FGF7 (Peprotech), 500 nM of A83-01 (Tocris Bioscience), 2% of B27 (Thermofisher), and 10 μM of Y-27632 ROCK inhibitor (Abmole Bioscience). Over the subsequent 2-3 weeks, the mouse bladder organoids were either passaged every 5 days or sheared with a fire polished glass pipette to reduce the size of the organoids. Thereafter, the organoids were either used immediately for infection experiments or cryopreserved in freezing media (60% FBS, 30% Advanced DMEM/F-12, 10% DMSO) at −180°C for subsequent experiments.

### UPEC culture and injection of mouse bladder organoids

Uropathogenic *Escherichia coli* (UPEC) strain CFT073 was isolated from a pyelonephritis patient (provided by H.L.T. Mobley). A derivative strain expressing yellow fluorescent protein (YFP) was generated by electroporation of CFT073 with the episomal plasmid PZA32-YFP. To induce expression of type 1 pili, UPEC was grown in LB media containing 25 μg/mL chloramphenicol in non-shaking condition at 37°C for 2 days prior to the experiment.

Cryopreserved tdTomato mouse bladder organoids were thawed and recultured in BME for five days before the experiment. On the day prior to the experiment, the organoids were washed twice with ice-cold DMEM (without antibacterials or antifungals) followed by centrifugation at 100 g for 5 minutes to remove the spent BME. They were then seeded in fresh BME inside a 35 mm ibidi μ-Dish in basal medium without antibacterial or antifungal supplementation. On the day of the experiment, stationary-phase UPEC were microinjected into the organoid lumen using a Pneumatic PicoPump (WPI). The micropipettes used for microinjection were prepared from thin wall glass capillary (TW100F-4 with length 100 mm and diameter 1mm) using a Flaming/Brown Micropipette Puller (Sutter Instruments model P-87) set at pressure 360, heat 866, Vel 200. Prior to the microinjection of organoids with UPEC, the micropipettes were filled with 10 μL of 1:1 volume dilution of UPEC culture with a Phenol Red solution (Sigma-Aldrich) using Microloader flexible tips (Eppendorf). This enabled the user to visualize the injected volume. The micropipettes were then cut with a sharp scalpel under a stereomicroscope (Olympus SZX-16) to generate a wider tip. The tip size of the cut micropipette was verified by setting pressure conditions adjusting the input pressure on PicoPump to eject ca. 1 nL volume of liquid in mineral oil on a thin film of paraffin on a Zeiss coverslide (corresponding to 100 μm). We verified the inoculum size by plating for colony forming units on LB agar. Typically, we aimed to inject medium-sized organoids (100-300 μm in diameter) with a clearly distinguishable lumen as these were not only the easiest organoids to inject but were also of a size that was amenable to whole organoid live-cell imaging.

For the experiments where the infected organoids were co-cultured with mouse bone marrow derived neutrophils, the injected organoids were then removed from fresh BME by pipetting 2 mL of ice-cold Cell Recovery Solution (Corning) after removing MBM from ibidi μ-Dish. The BME hydrogel was mechanically dissociated with a P1000 pipette and the resulting liquid gel was placed inside a 15 mL falcon tube pre-coated with a 1% BSA solution. The ibidi μ-Dish was washed twice with an additional 1.5 mL of Cell Recovery Solution to collect the organoids, and the resulting 5 mL of Cell Recovery Solution containing organoids inside the 15 mL falcon tube was further mechanically dissociated with a non-tapered glass pipette pre-coated with a 1% BSA solution to ensure the complete removal of the organoids from the BME hydrogel. The cell recovery solution containing infected organoids was kept on ice for 30 minutes to completely liquefy the BME. The cell recovery solution was then exchanged through two washes with 10 mL of basal medium (supplemented with 10% FBS) and the organoids were centrifuged at 100 g for 5 minutes at 4°C to remove the recovery solution. The organoid pellet was then resuspended in a collagen gel that was suitable to observe the migratory dynamics of the neutrophils.

### Isolation of neutrophils and labelling with CellTracker™ dye

Neutrophils were isolated from bone marrow of three female WT mice aged between 2 and 4 months (Charles River) per experiment. Briefly, mice were euthanized by CO_2_ overdose and the femur and tibia were isolated. The isolated bone marrow was crushed with a mortar and pestle and resuspended in 5 mL of cold SM++ solution (HBSS + 2% serum + 25 mM HEPES). The tissue homogenate was then passed through a pre-wet 40 μm filter. An additional volume of cold SM++ solution was added to the filtered cell suspension to make a total volume of 50 mL and then pelleted down at 300 g for 5 minutes. The cell suspension was then processed in batches of 10^7^ cells and neutrophils were isolated using positive selection with the Miltenyi Anti-Ly6G kit with a high degree of purity using the protocol described by the supplier (Swamydas et al., 2015). Isolated Ly6G mice neutrophils were stained with CellTracker™ Deep Red (Thermofisher) in a serum-free RPMI phenol red free medium at 1 μM and incubated for 30 minutes in 5 mL cell suspension inside 50 mL falcon tube maintained at 37°C. Neutrophils were labelled with Post-CellTracker™ and subsequently washed twice with 10 mL of 20% FBS in RPMI phenol red free medium to remove the unbound dye. The labelled neutrophils were kept briefly at room temperature before introduction into a co-culture with infected organoids inside the collagen gel.

### Co-culture of mouse neutrophils and infected organoids in collagen gels

Collagen gels were used to co-culture infected organoids and mouse neutrophils. The collagen gel master mix buffered at pH 7.0 was made by adding 312 μL of ice-cold native bovine collagen with 4 μL of 1M HEPES, 4 μL of 1M sodium bicarbonate, 40 μL of 1X DMEM/F-12, and 40 μL of 10X DMEM/F-12. The collagen gel mixture was stored on ice before use and care was taken to avoid generating bubbles when pipetting. Freshly isolated neutrophils (ca. 10^7^cells labeled with CellTracker™ were resuspended in 88 μL of a premade mixture of collagen gel master mix (10X DMEM/F-12, 1X DMEM/F-12, HEPES, sodium bicarbonate) without native bovine collagen on ice and then mixed with infected organoids. 156 μL of collagen solution was added twice to the suspension of infected organoids and neutrophils on ice followed by rapid mixing with a P200 pipette. This collagen gel suspension of infected organoids and neutrophils was pipetted on surface of an ibidi μ-Dish, which was pre-treated with an exposure to plasma (Diener, pressure 60 and time 60 s) to increase the hydrophilicity of the surface. This enabled a uniform thin layer (1-2 mm) of collagen gel to be deposited on the dish which was required for imaging with a confocal microscope. The collagen gel was then incubated at 37°C and 5% CO_2_ for 30 minutes to allow the collagen gel to polymerize and solidify. Lastly, 1.5 mL of a 25 ng/mL solution of murine Granulocyte Colony Stimulating Factor (G-CSF) was added to the co-culture of infected organoids and neutrophils to ensure that the neutrophils were exposed to chemokines that stimulate maturation (Ingersoll et al., 2008; Semerad et al., 2002).

#### Quantitative real-time PCR (qRT-PCR) of infected organoid samples

Organoids were infected as described above. 4 hours after injection of UPEC, the cell culture media surrounding the organoids was removed, the organoids were incubated with the appropriate volume of RNA lysis buffer (RNAeasy Plus Micro Kit, Qiagen), RNA was isolated as per the manufacturer’s instructions and resuspended in 14 μl of DEPC-treated water. A similar procedure was followed for uninfected controls. 11 μl of the RNA-containing solution was then used to generate cDNA using the SuperScript®IV First-Strand Synthesis System with random hexamers (Invitrogen), which was stored at −20 °C. Specific primers for used for the qRT-PCR are listed in Table S2. qRT-PCR reactions were prepared with SYBR®Green PCR Master Mix (Applied Biosystems) with 500 nM primers, and 1 μl cDNA. Reactions were run as absolute quantification on ABI PRISM®7900HT Sequence Detection System (Applied Biosystems). Amplicon specificity was confirmed by melting-curve analysis.

### Immunofluorescence of uninfected and infected mouse bladder organoids

For immunofluorescence imaging of paraffin-embedded slices, cryopreserved mouse bladder organoids were thawed and maintained in BME culture for five days before the experiment. Both uninfected and infected organoids were fixed with 4% paraformaldehyde (PFA) for 6 hours at room temperature with occasional mechanical dissociation with a P1000 pipette. Fixed organoids were then washed twice with PBS and centrifuged at 100 g for 5 minutes in a 15 mL falcon tube pre-coated with 1% BSA. The organoids then resuspended in 50 μL of prewarmed Histogel (Thermofisher) at 50 °C and pipetted out as a small hemispherical dome inside a 1 cm Tissue-Tek Cryomold. The cryomold was placed on a cold ice plate for solidification. Subsequently, the hemispherical Histogel was processed for paraffin embedding. Organoids embedded in paraffin were cut into 4 μm slices. The thin paraffin sections were deparaffinized and rehydrated by immersing the slides through the following solutions: xylene, three washes for 5 minutes each; 100% ethanol, two washes for 10 minutes each; 95% ethanol, two washes for 10 minutes each; 70% ethanol, two washes for 10 minutes each; 50% ethanol, two washes for 10 minutes each; PBS, three washes for 5 minutes each. Rehydrated slides were then processed for heat induced antigen retrieval using 10 mM citrate buffer, pH =6.0. Slides were washed with 1X PBS, permeabilized with 0.15 % Triton X-100 for 15 minutes, washed twice with 1X PBS, and blocked with 1% BSA in PBS for 1 hour. The boundaries of paraffin sections were marked with a hydrophobic pen and slides were labelled with a permanent ethanol-resistant marker. Incubation with primary antibodies at a dilution of either 10 μg/mL or 1:100 was performed overnight in an antibody incubation buffer comprising 1% BSA and 0.01% Triton-100 in PBS. Slides were subsequently washed three times with PBS for 10 minutes each. Incubation with secondary antibodies was performed at 2 μg/mL in antibody incubation buffer for 1 hour at room temperature. In each step, excess antibody was removed by washing three times with PBS for 10 minutes each. Cell nuclei were stained with DAPI (5 μg/mL) for 30 minutes and slides were mounted with Fluoromount-G™ mounting medium (Thermofisher) overnight in a dark chamber. For immunofluorescence imaging of whole organoids, uninfected and infected organoids were cultured for 6 hours in collagen gels on Ibidi μ-Dishes, fixed with 1 ml of 4% PFA for 1 hour, and antibody labelling was performed as described above.

### Time-lapse confocal imaging of infected organoids

All imaging experiments were conducted using a Leica SP8 confocal microscope in the inverted configuration with a temperature-controlled microscope environmental chamber maintained at 37°C and a stage-top chamber that allowed supplementation with 5% CO_2_ (OKOlabs). Time-lapse imaging was conducted using a Leica HC FLUOTAR 25X (NA 0.95) multi-immersion objective. To maintain the water immersion for the objective water was pumped to the ring around the water objective at 9 Hz with pumping duration 10 seconds and pumping interval 30 minutes, controlled by SRS software. Microinjected tdTomato-expressing organoids were identified and imaged on either two channels (for experiments which did not involve the addition of neutrophils) or three channels (for experiments where neutrophils were added), in each case the multiple channels were acquired during the same imaging sequence to improve the temporal resolution We therefore chose laser excitation wavelengths of 500 nm (YFP), 555 nm (tdTomato), and 630 nm (CellTracker™ Deep Red) to enable multi-channel imaging within the same sequence with reduced spectral overlap. Images were acquired with a scan speed of 700 Hz and a zoom factor of 2.25 which provided an XY resolution of between 200-450 nm, corresponding to images that were 1024 × 1024 pixels or 512 × 512 pixels respectively. Z-stacks were acquired with 0.5 or 1 μm step sizes.

### Additional experimental steps for experiments involving ampicillin treatment

After UPEC microinjection, MBOs were resuspended in collagen and supplemented with 1.5 mL of MBM medium following collagen polymerization for 30 minutes. Prior to the experiment, Ibidi μ-Dishes were modified by inserting bent metallic tubes (1.00/0.75 X 20mm, Unimed Catalogue 200.010-A) through the lid and sealing these devices with PDMS cured in an oven at 80°C for 1 hour. Flexible 0.76 mm × 1.65 mm × 1 mm tubing (Mono-Lumen Freudenberg Medical) was connected to these tubes to facilitate medium addition and removal. A setup period of 2 to 3 hours was typically required and is not considered within the time duration of the experiment. Once imaging commenced, bacteria were allowed to grow for a 3-hour “growth phase” (0-165 minutes). The medium in the μ-Dish was then replaced with fresh medium containing ampicillin at 64.5 μg/mL, corresponding to 10X-MIC. The dynamics of bacterial growth and killing in the presence of ampicillin was then monitored for a 3-hour “treatment phase” (ca. 165-345 minutes), after which ampicillin was removed from the extracellular medium by gentle exchange with fresh MBM media using a 20 mL syringe. Bacterial growth after ampicillin washout was then monitored for a further 3-hour “regrowth phase” (345-525 minutes). The drift in focus was adjusted manually at frequent intervals throughout the experiment.

For the experiments to test the kinetics of ampicillin diffusion into the organoids, Organoids were ‘mock-infected’ by injection with pHrodo™ Deep Red E. coli BioParticles™, diluted ten times in LB (to closely mimic the injection with UPEC). The injected organoids were then resuspended in collagen in the μ-Dish as described previously. The pHrodo particle injected organoids and uninjected bystander organoids were imaged for a period of 45 minutes with an imaging period of 15 minutes. Thereafter, we administered a 0.2 μM Lucifer Yellow (reconstituted in a 0.1M solution of Tris buffer at pH 7.4) along with ampicillin (at 64.5 μg/mL) in MBM medium. The medium containing Lucifer Yellow and ampicillin was removed at the end of 3-hour treatment period from the extracellular medium by gentle exchange with fresh MBM media using a 20 mL syringe. The mean intensity of Lucifer Yellow was computed within injected and bystander organoids by creating intraorganoid mask using tdTomato fluorescence of the epithelial cells as described in the section below. Mean Lucifer intensity in the extraorganoid environment was calculated within a bounding box (20 × 20 × 33 μm^3^) located within an appropriate volume outside the organoid.

### Image analysis for confocal live-cell imaging

Image analysis was performed with Bitplane Imaris 9.5.1. and is outlined in greater detail in Fig. S4 (for the analyses related to Fig. 2) and Fig. S7 (for the analyses related to Fig. 3 and 4). The following is a step-by-step summary of the analysis pipeline as applied for the analysis in Fig. 3 and 4. The time-lapse images acquired have four optical channels for UPEC (first channel), uroepithelial cells (second channel), neutrophils (third channel), and transmitted light (fourth channel).

First, a background subtraction for the neutrophil channel was performed due to spectral overlap between the tdTomato expression from the epithelial cells and the CellTracker Deep Red staining of the neutrophils. This procedure allowed for a more robust detection of neutrophils inside the organoids. This created a fifth channel (Fig. S7B). Next the organoid surfaces were generated using the ‘Surface Generation Tool’ as outlined in Fig. S7C. When required, adjacent bystander organoids were separated by setting the expected volume of the organoids using the ‘seed point diameter’ function to automatically segment the organoids (depending upon the size of nearby organoids). In some cases, adjoining organoid surfaces were orthogonally trimmed with the ‘cut’ tool. The organoid surface was then used as a mask to spatially distinguish intraorganoid vs. extraorganoid bacterial and neutrophil populations (Fig. S7D,E). These were stored as separate imaging channels in Imaris. Lastly, the intraorganoid and extraorganouid neutrophil channels were segmented to generated structures whose surfaces which are depicted in the processed images in Fig. 3A-C, 4A-C, S9A-C and whose volumes are quantified in Fig. 3D-F, S8A-C, S9D-I. In a similar fashion, the intraorganoid UPEC were segmented to generated structures whose surfaces are shown in Fig. 2B-I, S3A and whose volumes are used to calculate the intraoorganoid bacterial growth kinetics in Fig. 2J-M, S3C and total bacterial volume inside the organoid in Fig. 4D, S8A-C, S9G-I.

#### Neutrophil spot detection

The masked sixth channel was also used to detect neutrophil spots (Size, 8 μm; Quality, 4 to 8) inside the organoid. Neutrophil spots in the field of view were detected on the neutrophil channel (third channel) with parameters (Size, 8 μm; Quality, 4 to 8). Neither neutrophil spots nor neutrophil surfaces were tracked over time due to the technical difficulties with imaging speed and amorphous shapes of migrating neutrophils around infected organoids. Average neutrophil sphericity (≥0.5) for neutrophils around an organoid was obtained with time-collapsed information of average neutrophil sphericity for all the neutrophil surfaces per time point. In cases of bigger neutrophil clusters, average neutrophil sphericity was obtained by excluding sphericity of the bigger neutrophil cluster (<0.5). For UPEC volume inside the organoids, the first channel was sufficient to create the surface (Threshold, 15; Smooth Surfaces Detail, 0.5 or 1 μm)).

### Serial block face-scanning electron microscopy (SBEM) of an infected organoid

tdTomato organoids expressing tdTomato were grown in Ibidi μ-Dishes, microinjected with UPEC, which was allowed to grow within the organoid lumen for six hours. After two hours of bacterial growth, injected organoids were removed from fresh BME, resuspended in a collagen gel surrounded by isolated bone marrow derived neutrophils as described above. The organoids were then plated in MBM supplemented with 25 ng/mL of mouse G-CSF on a plasma-treated MatTek dish with a gridded coverslip that enabled correlative light and electron microscopy. After 6 hours, organoids were fixed for 1 hour at room temperature in a 15 mL falcon tube containing a mix of 1% glutaraldehyde and 2% paraformaldehyde in 0.1 M phosphate buffer (pH 7.4). Infected organoids were fixed for one hour and then washed twice with PBS. The organoids were then screened using optical microscopy to rapidly identify infected organoids with at least one IBC and neutrophils within the lumen suitable for volumetric electron microscopy. The selected organoid was then imaged on the Leica SO8 microscope using a 10X objective to acquire a fluorescent image, which was used to identify and locate the target organoid on the gridded coverslip for correlative light and SBEM. This fixed organoid was kept overnight at 4°C in the fixative.

The fixed organoids were then postfixed in potassium ferrocyanide (1.5%) and osmium (2%), then stained with thiocarbohydrazide (1%) followed by osmium tetroxide (2%) alone. They were finally stained overnight in uranyl acetate (1%) and washed in distilled water at 50°C before staining with lead aspartate at a pH of 5.5 at the same temperature. The entire cover slip, with the organoids attached, were dehydrated in increasing concentrations of alcohol and then embedded in durcupan resin and hardened at 65°C for 24 hours. The total thickness of the coverslip and resin was minimized to around 1 mm. Once hardened, the coverslip was removed, and the region of resin containing the organoid of interest was cut away from the others with a scalpel blade. This piece was glued, with conductive glue, to an aluminium holder, and then placed inside the scanning electron microscope (Merlin, Zeiss NTS) integrated with an in-chamber ultramicrotome device (3View, Gatan). Serial images,100 nm part, were collected from the block face using a beam energy of 1.7 kV and 350 pA of current. Each image contained 6144 × 4608 pixels with a pixel size of 30 nm, and the stack contained 960 serial images. The overall volume imaged was 184.32 × 138.24 × 96 μm^3^.

### Labelling of bacteria, epithelial cells, neutrophils, and organoid lumen in SBEM images

Identification and labelling of different features (e.g., lumen boundary, neutrophils, bacteria) was done manually; features whose identity was doubtful were not included in the analysis. All the different bacterial categories were identified on each slice of the full stack (960 slices). We identified five different categories of bacteria based on their location: extracellular bacteria in the organoid lumen, bacteria within neutrophils, bacteria within an IBC, bacteria within the cytoplasm of a uroepithelial cell, and pericellular bacteria sandwiched between the boundaries of adjacent cells. The injection site could be identified by the asymmetric distribution of lumenal bacteria with a higher abundance of extracellular lumenal bacteria towards one side of the lumen. We used the high fluorescence intensity of the large number of bacteria within the IBC to register and align the fluorescent image with the SBEM stack. In order to delineate structural features of the organoid such as the boundary of the organoid lumen, and the boundaries of the neutrophils as well as the overall organoid volume, a reduced stack (480 z-stacks) with X, Y, Z pixel resolution of 60 nm, 60 nm, and 100 nm was generated. We also used this reduced stack to mark the location of all the uroepithelial cell nuclei within the organoid. Neutrophils were identifiable in electron microscope images due to their multi-lobular nuclei and high electron density, and in cases where there was some ambiguity, the fluorescent images of neutrophils labeled with CellTracker™ Deep Red were used to verify neutrophil identity. Altogether we identified five neutrophils surrounding the organoid, and nine neutrophils were identified inside the organoid.

### 3D analysis and modelling of the infected organoid

Serial electron micrographs were imported into the TrakEM2 plugin in FIJI. They were then aligned and registered and the locations of all cells and bacteria annotated manually through the image stack. The organoid lumen was demarcated by segmenting the lumen boundary in each of the slices of the stack in each of the images in which it appeared. A similar procedure was also undertaken for the boundaries of the neutrophils and the cell boundary of the cells surrounding the IBC and containing the IBC. For ease of manipulating the large dataset, these reconstructions of the lumen and identification of neutrophils and uroepithelial cells was done on a reduced stack of 480 slices with corresponding pixel size of 60 nm and spacing between serial slices of 200 nm.

The segmentations of cells and the organoid lumen, and the coordinates of each mouse cell and bacterial cell, were then exported as 3D wavefront object in the TrakEM2 and imported into the Blender 3D modeling software. The “proximity tool” in the NeuroMorph toolset for Blender (Jorstad et al., 2018) was used to measure the distance of each bacterium to the wall of the lumen, as well as the organoid’s external surface. To visualize the organoid, with all bacteria, the particles system in the Blender software was used to place a single generic model of either a bacterium or a cell at the vertex that marked its 3D coordinate.

### Scanning electron microscopy of infected organoids

tdTomato-expressing organoids were infected with UPEC and maintained in a cell culture incubator for the subsequent 6 hours. After this period the organoids were fixed for 1 hour at room temperature in a 15 mL falcon tube containing a mix of 1% glutaraldehyde and 2% paraformaldehyde in 0.1 M phosphate buffer (pH 7.4). Fixed organoids were then seeded onto 12 mm poly-L-lysine-coated coverslips and stored overnight in the fixative. The organoids were then further fixed with 1% osmium tetroxide in 0.1 M cacodylate buffer, followed by a dehydration step in increasing concentrations of ethanol (50% to 100% in 10% increments, 10 minutes per incubation). The dehydrated organoids were then dried at the critical point of CO_2_ (Leica CPD300) before being ruptured with a 0.125 mm tungsten needle (FST, 10130-05) using a stereomicroscope (Leica M205). The ruptured organoid was then adhered to the surface of an aluminium stub using conductive tape (Election Microscopy Sciences, USA). This holder was coated with a 5 nm thick layer of gold palladium metal using a Q150 sputter coater (Quorum Ltd, UK), and then imaged in a scanning electron microscope (Merlin, Zeiss NTS). Images were acquired with an electron beam voltage of 1.50 kV and current of 80 pA with HE-SE2 detector.

