## Supplementary Material for "Early invasion of uropathogenic *Escherichia coli* into the bladder wall by solitary bacteria that are protected from antibiotics and neutrophil swarms in an organoid model"

**KEY RESOURCES TABLE**

| REAGENT or RESOURCE | SOURCE | IDENTIFIER |
| --- | --- | --- |
| **Antibodies** | | |
| Anti-LPS | Abcam | Cat#: ab 35654 |
| Anti-CK7 | Abcam | Cat#: ab209599 |
| Anti-CK8 | Abcam | Cat#: ab192468 |
| Anti-Ly6G -PE | Biolegend | Cat#: 127607 |
| Anti-CD11b -BV711 | Biolegend | Cat#: 101241 |
| Anti-flagellin | Abcam | Cat#: ab 93713 |
| Anti-p63 | Abcam | Cat#: ab 735 |
| Anti-Uroplakin3a | Santa Cruz | Cat#: sc-166808 |
| Donkey anti-Mouse IgG (H+L) Highly Cross-Adsorbed Secondary Antibody, Alexa Fluor 647 | Thermofisher | Cat#: A-31571 |
| Donkey anti-Rabbit IgG (H+L) Highly Cross-Adsorbed Secondary Antibody, Alexa Fluor 647 | Thermofisher | Cat#: A-31573 |
| Goat anti-Mouse IgG (H+L) Highly Cross-Adsorbed Secondary Antibody, Alexa Fluor 488 | Thermofisher | Cat#: A-11029 |
| **Bacterial and virus strains** | | |
| Uropathogenic Escherichia coli (UPEC) strain CFT073 | PMID: 2182540 | NCBI:txid199310 |
| CFT073-pZA32-YFP | This paper |  |
| **Biological samples** |  |  |
| **Chemicals, peptides, and recombinant proteins** | | |
| Native Collagen, Bovine dermis | AteloCell | Cat#: IAC-50 |
| B27 | Thermofisher | Cat#:17504044 |
| Human KGF/ FGF7 | Peprotech | Cat#:100-19 |
| Human FGF10 | Peprotech | Cat#:100-26 |
| Cultrex PathClear Reduced Growth Factor BME, Type 2 | Bio-Techne | Cat#: 3533-005-02 |
| Ampicillin | Sigma-Aldrich | Cat#: A9518-5G |
| Chloramphenicol | Sigma-Aldrich | Cat#:C1919-25G |
| Gibco^TM^ Trypsin-EDTA (0.05%), phenol red | Thermofisher | Cat#:25300054 |
| Gibco^TM^ Antibiotic-Antimycotic (100X) | Thermofisher | Cat#:15240062 |
| Gibco^TM^ Fetal Bovine Serum | Thermofisher | Cat#: 10270106 |
| Gibco^TM^ RPMI 1640 Medium, no phenol red | Thermofisher | Cat#:11835030 |
| RPMI-1640 medium | ATCC | Cat#:30-2001 |
| **Critical commercial assays** | | |
| Anti-Ly-6G MicroBead Kit, mouse | Miltenyi Biotec | Cat#: 130-092-332 |
| **Deposited data** | | |
| **Experimental models: cell lines** | | |
| **Experimental models: organisms/strains** | | |
| Gt(ROSA)26Sortm4(ACTB-tdTomato,-EGFP)Luo/J | Jackson Laboratory | Cat#: 007576; PMID: 17868096 |
| C57BL/6 Mouse | Charles River Laboratory | Cat#: C57BL/6NCrl |
| **Oligonucleotides** | | |
| **Recombinant DNA** | | |
| pZA32-YFP | PMID: 9092630 |  |
| **Software and algorithms** | | |
| Imaris 9.5.1 | Bitplane |  |
| **Other** | | |
| Advanced DMEM/F-12 | Thermofisher | Cat#: 12634010 |
| DMEM/F-12, no phenol red | Thermofisher | Cat#: 21041025 |
| HEPES, 100X | Thermofisher | Cat#:15630106 |
| Glutamax, 100X | Thermofisher | Cat#: 35050061 |
| A8301 | Tocris | Cat#: 2939 |
| Rho kinase inhibitor, Y-27632 | Abmole Bioscience | Cat#: 2939 |
| Glass Pasteur pipettes | VWR | Cat#: 612-1701 |
| Gas chamber for stages with k-frame insert (160x110mm) - magnetic model with sliding lid. | Okolab | Cat#: H201-K-FRAME |
| Flaming/Brown Micropipette Puller | Sutter Instruments | model P-87 |
| Capillary glass, 1.0 mm outer diameter, 0.75 mm inner diameter | WPI | TW100F-4 |
| 35mm holder - magnetic | Okolab | Cat#:1x35-M |
| µ-Dish 35 mm, high | Ibidi | Cat#:81156 |
| Flexible tubing, 0.76X1.65X0.45X15000 | Freudenberg Medical | Cat#:0045634143 |
| 1.00/0.75 x 20mm metallic tubes | Unimed | Cat#:200.010-A |
| Aladdin programmable pump | WPI | Cat#:PUMP-NE-1000 |
| Olympus SZX16 stereo microscope | Olympus |  |
| Leica SP8 confocal microscope | Leica |  |
| Olympus MVX10 stereo microscope | Olympus |  |
| CellTracker™ Deep Red Dye | Thermofisher | Cat#: C34565 |
| LB (Luria broth base, Miller′s modified) | Sigma-Aldrich | Cat#: L1900-1KG |
| Invitrogen™ DAPI | Thermofisher | Cat#: D1306 |
| TrypLE™ Express Enzyme (1X) | Thermofisher | Cat#: 12605010 |
| Phosphate Buffered Saline | Thermofisher | Cat#: 10010056 |
| Lucifer Yellow CH, Lithium Salt | Thermofisher | Cat#: L453 |
| pHrodo™ Deep Red E. coli BioParticles™ Conjugate for Phagocytosis | Thermofisher | Cat#: P35360 |

### Supplementary Movie Legends

**Supplementary movie 1**

**File Name: SMov1**, Description: **Intra-organoid UPEC growth is refractory to antibiotic clearance (Fig 2: B-E)**.

UPEC (in green) grows inside the organoid (organoid surface in magenta generated by Bitplane Imaris) for 3 hours during the growth phase (0-165 min). Bacterial growth is predominantly observed in the organoid lumen. Following addition of ampicillin at a concentration ten times the minimum inhibitory concentration (10 X MIC at ca. 165 min) in the extracellular medium, bacterial killing is observed. In response to ampicillin treatment (ca. 165-345 min), UPEC is observed to initially filament followed by lysis. After removal of the antibiotic (recovery phase ca. 345-525 min), bacterial regrowth appears to be restricted to the bladder epithelium or organoid wall.

**Supplementary movie 2**

**File Name: SMov2**, Description: **UPEC growth could be observed in the presence of the antibiotic (Fig 2: F-I)**.

UPEC (in green) grows inside the organoid (organoid surface in magenta generated by Bitplane Imaris) for 3 hours during the growth phase (0-165 min). Bacterial growth is predominantly observed in the organoid lumen. Following addition of ampicillin at a concentration ten times the minimum inhibitory concentration (10 X MIC at ca. 165 min) in the extracellular medium, bacterial killing is observed. In response to ampicillin treatment (ca. 165-345 min), UPEC is observed to initially filament followed by lysis. However, during the later stage of the ampicillin treatment (after 240 min), UPEC growth is observed in the bladder epithelium or organoid wall.

**Supplementary movie 3**

**File Name: SMov3**, Description: **Intra-organoid UPEC growth is refractory to antibiotic clearance (Fig S2**: **A1-A7**). An additional example showing growth of the bacteria within the lumen of an organoid during the growth phase (0-165 min), bacterial killing upon exposure to ampicillin treatment (ca. 165-345 min), and a slow regrowth after the removal of ampicillin (recovery phase ca. 345-525 min).

**Supplementary movie 4**

**File Name: SMov4**, Description: **Animated Z stack and 3D rotations highlight the presence of solitary bacteria in the organoid wall at the end of a treatment with a 10X MIC solution of ampicillin (Fig S5**: **A-B**). UPEC was stained with anti-LPS antibody (shown in green) and umbrella-like cells lining the lumen were identified with anti-CK8 staining (shown in amber). Epithelial cells expressing the red-fluorescent protein tdTomato within cell membranes are shown in magenta.

**Supplementary movie 5**

**File Name: SMov5**, Description: **Animated Z stack and 3D rotations highlight the presence of solitary bacteria in the organoid wall at the end of a treatment with a 10X MIC solution of ampicillin) (Fig S5**: **C-D**). UPEC was stained with anti-LPS antibody (shown in green) and umbrella-like cells were identified with anti-CK8 staining (shown in amber). Epithelial cells expressing the red-fluorescent protein tdTomato within cell membranes are shown in magenta.

**Supplementary movie 6**

**File Name: SMov6**, Description: **Neutrophil form a persistent swarm in response to bacterial infection inside the organoid (Fig 3**: **A1-A5**).

Surfaces of intra-organoid neutrophils (amber), extra-organoid neutrophils (blue) and infected (magenta) and uninfected (red) organoids were generated using a Bitplane Imaris analysis pipeline. UPEC (green) are shown without image processing to identify individual bacteria. During the course of infection, UPEC grows inside the organoid (observed by increase in green fluorescence), and neutrophils migrate into the lumen of an infected organoid (but not the adjacent uninfected organoid (shown in red). A neutrophil swarm is formed around the bacteria in the lumen and persists over the course of the experiment. The neutrophil swarm kills a majority of the bacteria in the organoid lumen.

**Supplementary movie 7**

**File Name: SMov7**, Description: **Neutrophil form a transient swarm in response to bacterial infection inside the organoid (Fig 4**: **B1-B5**).

A transient neutrophil swarm (amber) is formed around the bacteria in the organoid. Following the bacterial killing, the neutrophil cluster disaggregates, and neutrophils migrate back out of the organoid to the surrounding collagen matrix.

**Supplementary movie 8**

**File Name: SMov8**, Description: **Neutrophil form a dynamic swarm in response to bacterial infection inside the organoid (Fig 4**:**C1-C5)**.

A dynamic neutrophil swarm (amber) is formed around the bacteria in the organoid. Neutrophils volume inside the organoid is observed to fluctuate during the course of the experiment.

**Supplementary movie 9**

**File Name: SMov9**, Description: **Neutrophil form a persistent swarm in response to bacterial infection inside the organoid (Fig S5**:**A1-A5)**.

Another example of persistent neutrophil swarm formed inside the infected organoid.

**Supplementary movie 10**

**File Name: SMov10**, Description: **Neutrophil form a transient swarm in response to bacterial infection inside the organoid (Fig S5**:**B1-B5**).

Another example of transient neutrophil swarm formed inside the infected organoid.

**Supplementary movie** **11**

**File Name: SMov11**, Description: **Neutrophil form a dynamic swarm in response to bacterial infection inside the organoid (Fig S5**:**C1-C5**).

Another example of dynamic neutrophil swarm formed inside the infected organoid.

**Supplementary movie 12**

**File Name: SMov12**, Description: **Intracellular bacterial community is protected from surrounding neutrophils (Fig 4**: **A1-A5**).

Bacteria inside the IBC (imaging time from start of the experiment: 448 min) are protected from a persistent swarm inside the infected organoid. Following the event of IBC shedding (relative time: 0-60 min, imaging time: 448-496 min), bacteria are killed by the surrounding neutrophils All the neutrophils (inside and outside the organoid) are shown in amber. Segments during the time period (absolute time:192-448 min) were out of focus and could not be captured during imaging. All the neutrophils (inside and outside the organoid) are shown in amber.

**Supplementary movie 13**

**File Name: SMov13**, Description: **Neutrophil swarms spread the IBC bacteria (Fig 4**: **B1-B5**).

Bacteria inside the IBC (imaging time from start of the experiment: 160-240 min) are protected from a persistent swarm inside the infected organoid. Following the event of IBC shedding (relative time: 0-32 min, imaging time: 240-272 min), majority of the bacteria are killed by the surrounding neutrophil swarm. However, some of the bacteria are spread away by the persistent swarm. Solitary bacterial in the organoid wall did not originate from an IBC. These results demonstrate additional niches within the layers of the bladder epithelium beyond IBCs that can also harbour subpopulations of bacteria that are resistant to clearance by antibiotics or neutrophils. Neutrophils outside the organoid are not shown for clarity.

**Supplementary movie 14**

**File Name: SMov14**, Description: **Volumetric electron microscopy reveals five distinct bacterial niches within an infected organoid (Fig 5**: **F-I**).

Model derived from the serial electron microscopy images of the entire organoid in which all bacteria and cells were plotted. Video shows the interior of the organoid, revealing the lumen (brown), and epithelial cells (grey), as well as the neutrophils (cyan). Bacteria located within the five niches corresponding to are colored violet (lumenal), grey (neutrophil), lemon yellow (IBC), red (pericellular), and green (intracellular) (Please note the change in colors used for labelling pericellular bacteria, intracellular bacteria and neutrophils between the **supplementary movie 12** and **Fig 5: F-I**). An IBC is seen surrounded by neutrophils. Solitary intracellular and pericellular bacterial sub-populations are visible around all the quadrants of the whole organoid. The width of this organoid is 85 µm.

### Supplementary Figures and Legends

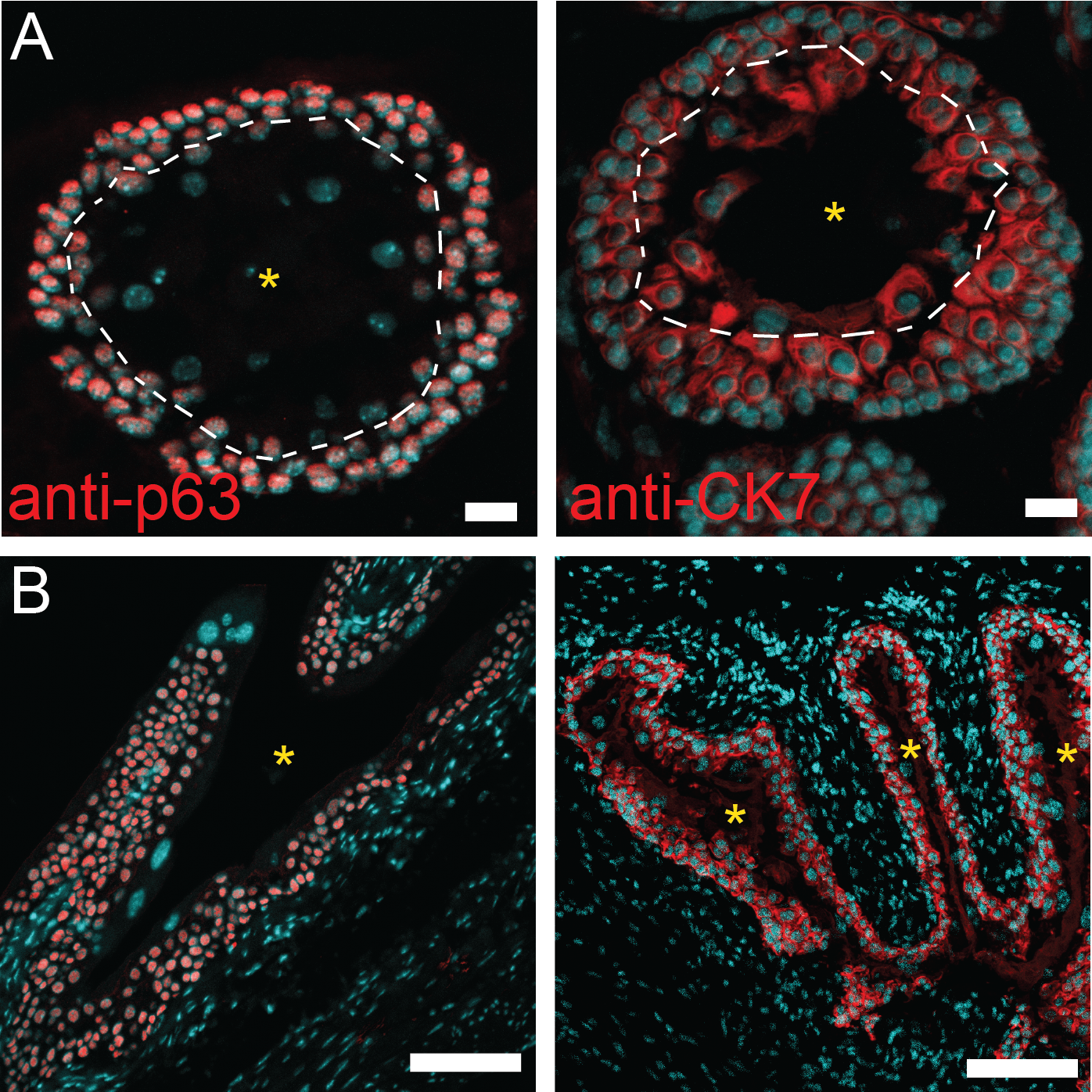

**Figure S1:** Mouse bladder organoids recapitulate the stratification of bladder uroepithelium. Intermediate and basal layers of the uroepithelium were identified in mouse bladder organoids (**A**) or explanted mouse bladder tissue (**B**) by immunofluorescence staining with anti-p63 (left panels) or anti-cytokeratin 7 (anti-CK7, right panels) antibodies. Cell nuclei were labeled with DAPI (cyan). The lumens in the bladder organoid and mouse bladder slices are indicated with yellow asterisks. The dashed white lines demarcate the boundary between the umbrella-like cell and intermediate cell layers. Scale bars: 20 μm in (**A**), 100 μm in (**B**).

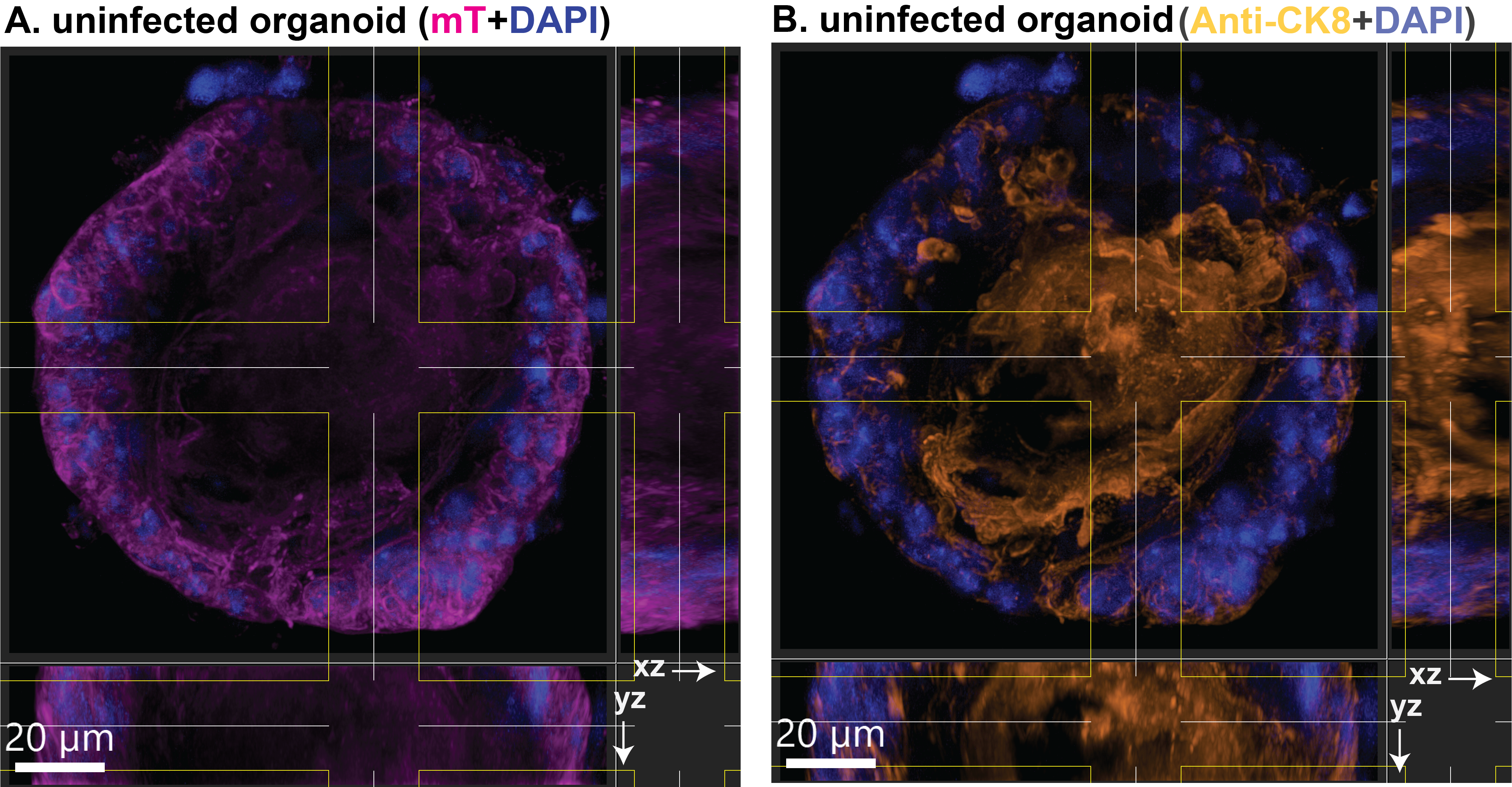

**Figure S2.** (**A, B**) Extended orthogonal section views of a 3D volume (20 x 20 x 20 μm^3^) provide an overview of biomarkers expressed at the boundary of the lumen of an uninfected organoid. The main image, XZ, and YZ panels show a maximum intensity projection within this volume along the Z, Y, and X axes, respectively. (**A**) Labelling of all cells in the organoid by td-Tomato expression (magenta) surrounding the organoid lumen in the center. (**B**) Immunostaining with anti-CK8 antibody shows the presence of umbrella-like cells within this volume.

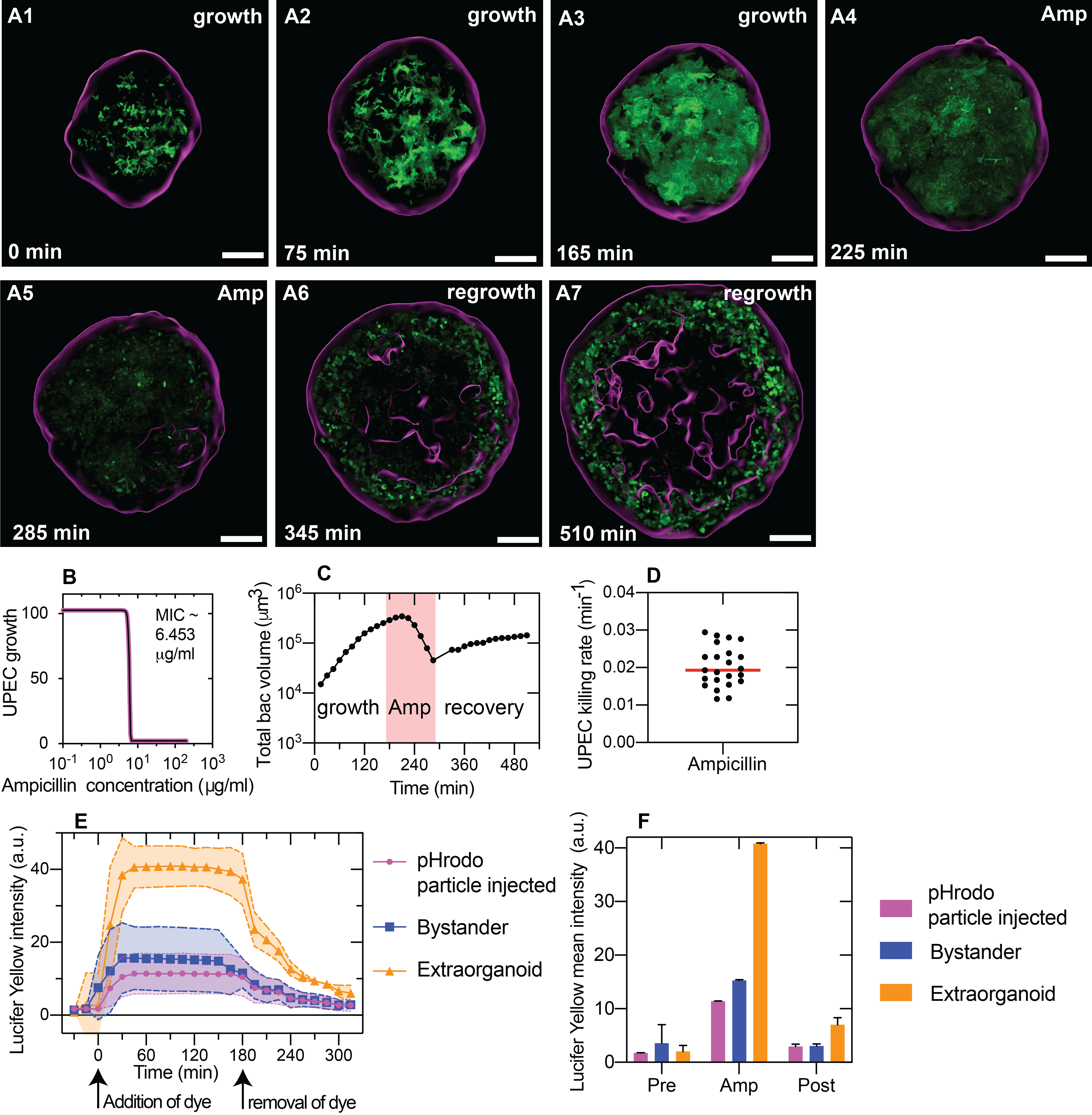

**Figure S3**. (**A1-A7**) An additional example showing bacterial growth within the organoid lumen, killing upon exposure to ampicillin, and slow regrowth within the organoid wall after ampicillin washout. (**B**) Corresponding time profiles for intra-organoid bacterial volume for the organoid in panels (**A1-A7**). (**C**) Measurement of ampicillin minimum inhibitory concentration (MIC) in mouse bladder organoid medium (DMEM) for the UPEC strain used in these experiments. (**D**) Scatter plot of the rate of decline of intra-organoid bacterial volume (“killing rate”) during ampicillin treatment, calculated by fitting the portion of the curves in Fig. 3M, after the maximum volume attained, with a linear fit. (**E**) Plot of mean intensity of lucifer yellow within a 512 x 512 x 33 μm^3^ volume within extraorganoid environment (n = 29), bystander organoids (n = 7), and pHrodo particle injected organoids (n = 22). Lines depict mean values, and the shaded regions depict the standard deviations. (**F**) Bar plots of the mean intensity across the three conditions in (E) before, during, and after removal of the dye from the medium surrounding the organoids. The mean intensity of Lucifer Yellow was calculated from three time points before (-45 minutes to 0 minutes), during (90-120 minutes), and after removal of ampicillin dye (300-330 minutes). Scale bars, 50 μm in (**A1-A7**).

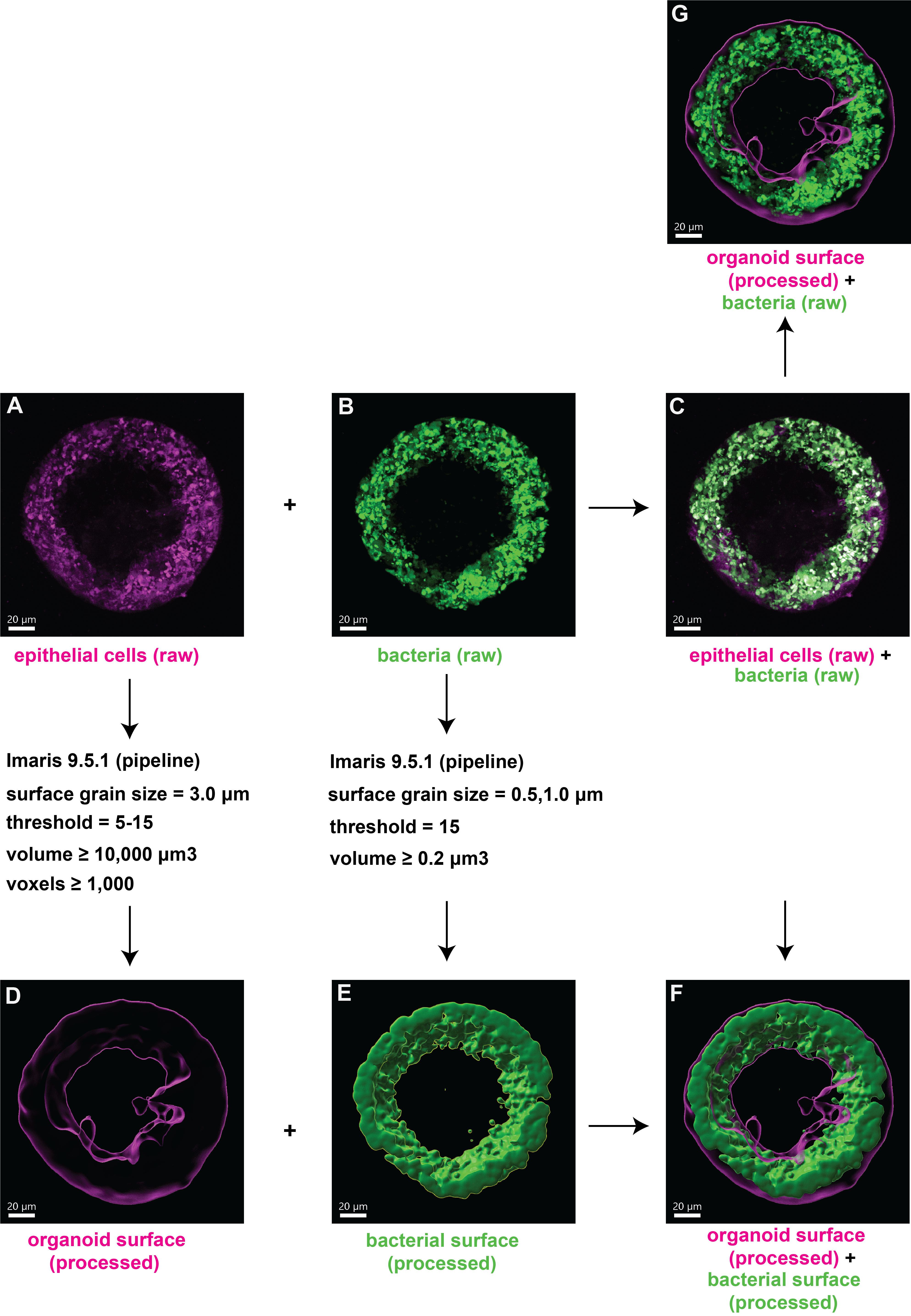

**Figure S4:** An illustrated example of the image processing pipeline for Figure 2 and Figure S3. (**A**, **B**) Raw data for tdTomato expression at the junctions of epithelial cells and YFP expression by UPEC, respectively. The raw images are processed using the ‘Surface Generation’ tool in Imaris with the given parameters to obtain the processed images in (**D**) and (**E**). Examples of overlays of the two channels using raw data in both channels (**C**), processed data in the organoid channel but raw data in the bacterial channel (**G**), or processed data in both channels (**F**).

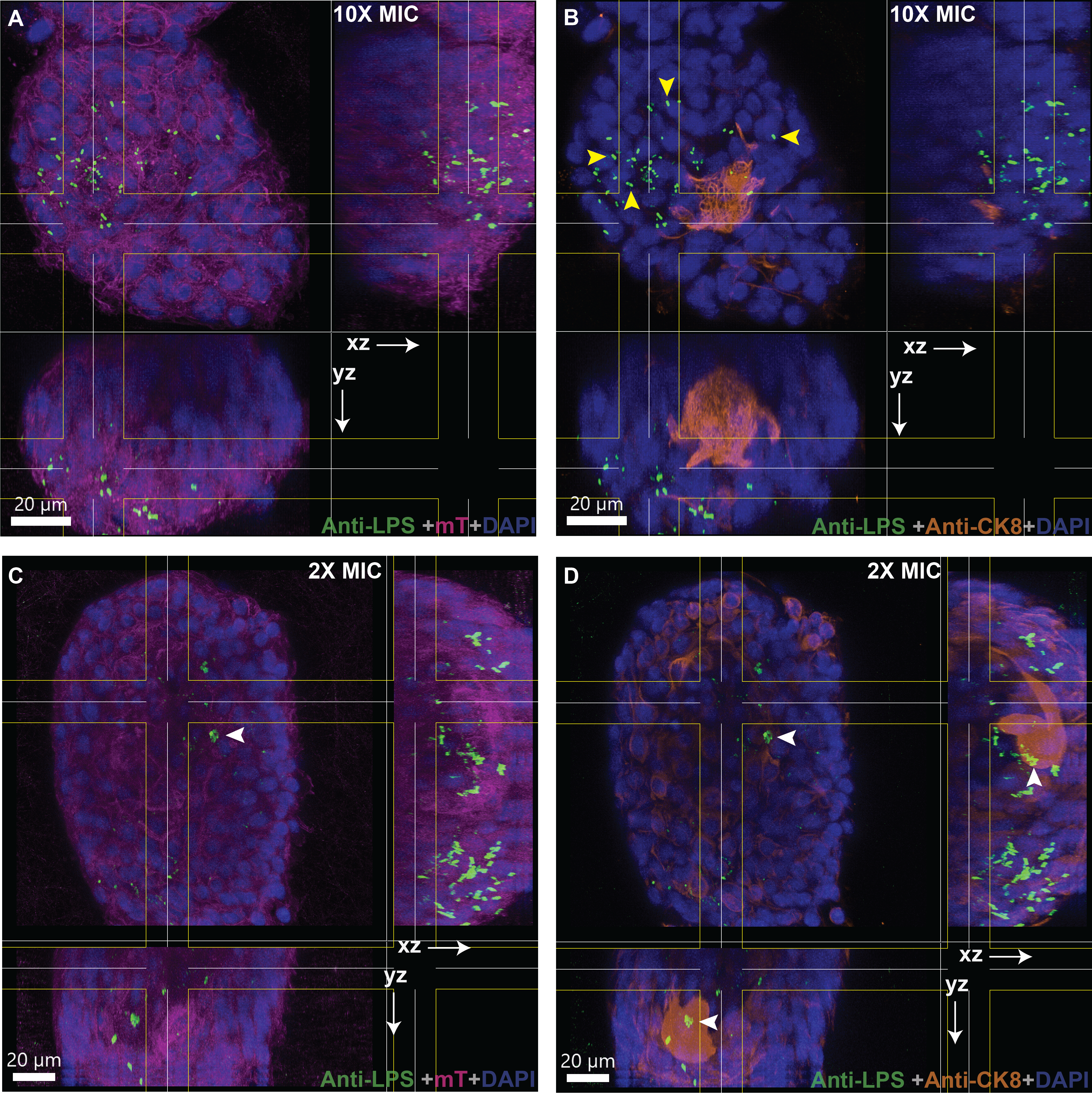

**Figure S5:** Orthogonal section views of a 3D volume (20 x 20 x 20 μm^3^) within the organoid wall of infected organoids after ampicillin treatment at 10X MIC (**A**, **B**) or 2X MIC (**C**, **D**). The main image, XZ, and YZ panels show a maximum intensity projection within this volume along the Z, Y, and X axes, respectively. UPEC are identified via an anti-LPS antibody (spring green). mT labelling of the uroepithelial cells is shown in magenta in (**A**, **C**) and cytokeratin 8 staining is shown in amber in (**B**, **D**). Examples of solitary bacteria in the bladder wall (CK8- cells) are indicated by yellow arrows in (**B**). An example of a bacterial clump within CK8+ cells is indicated by a white arrowhead in (**C**, **D**).

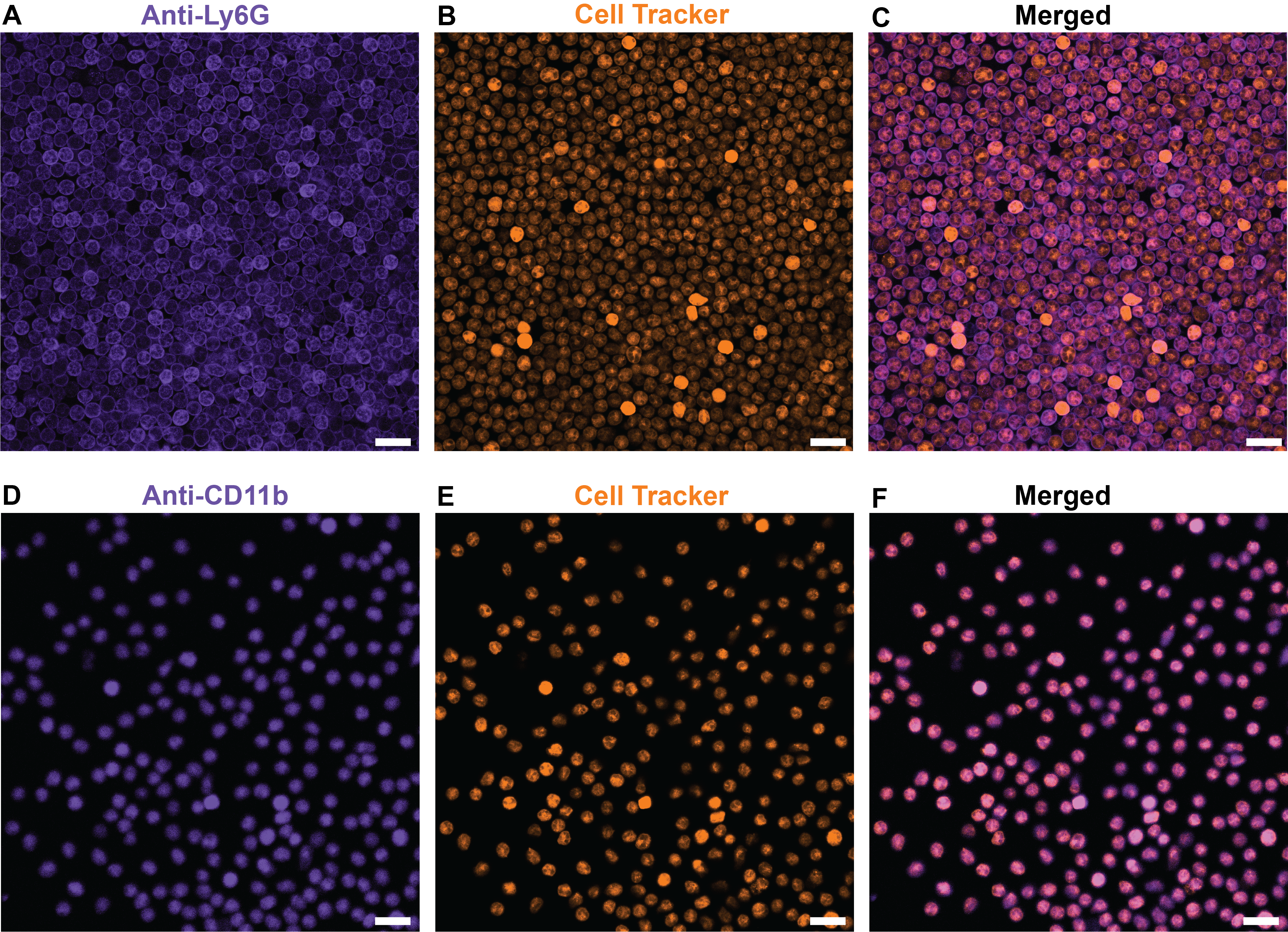

**Figure S6:** Verification of the neutrophil isolation protocol. Isolated neutrophils are labelled with CellTracker Deep Red (B, E) and immunostained for Ly6G (**A**) and CD11b (**D**). Merged images in each case (**C**, **F**) show a high degree of isolation purity.

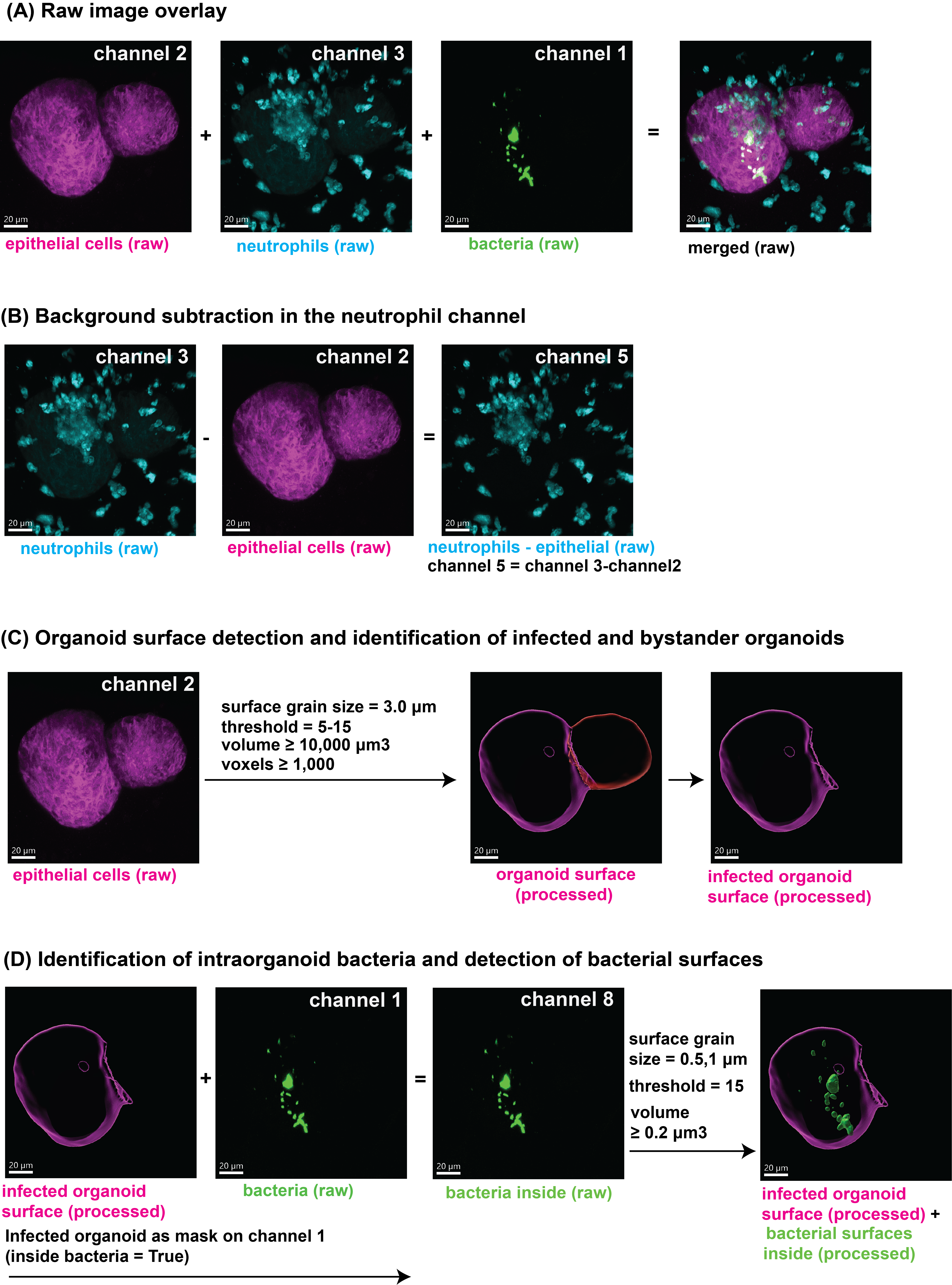

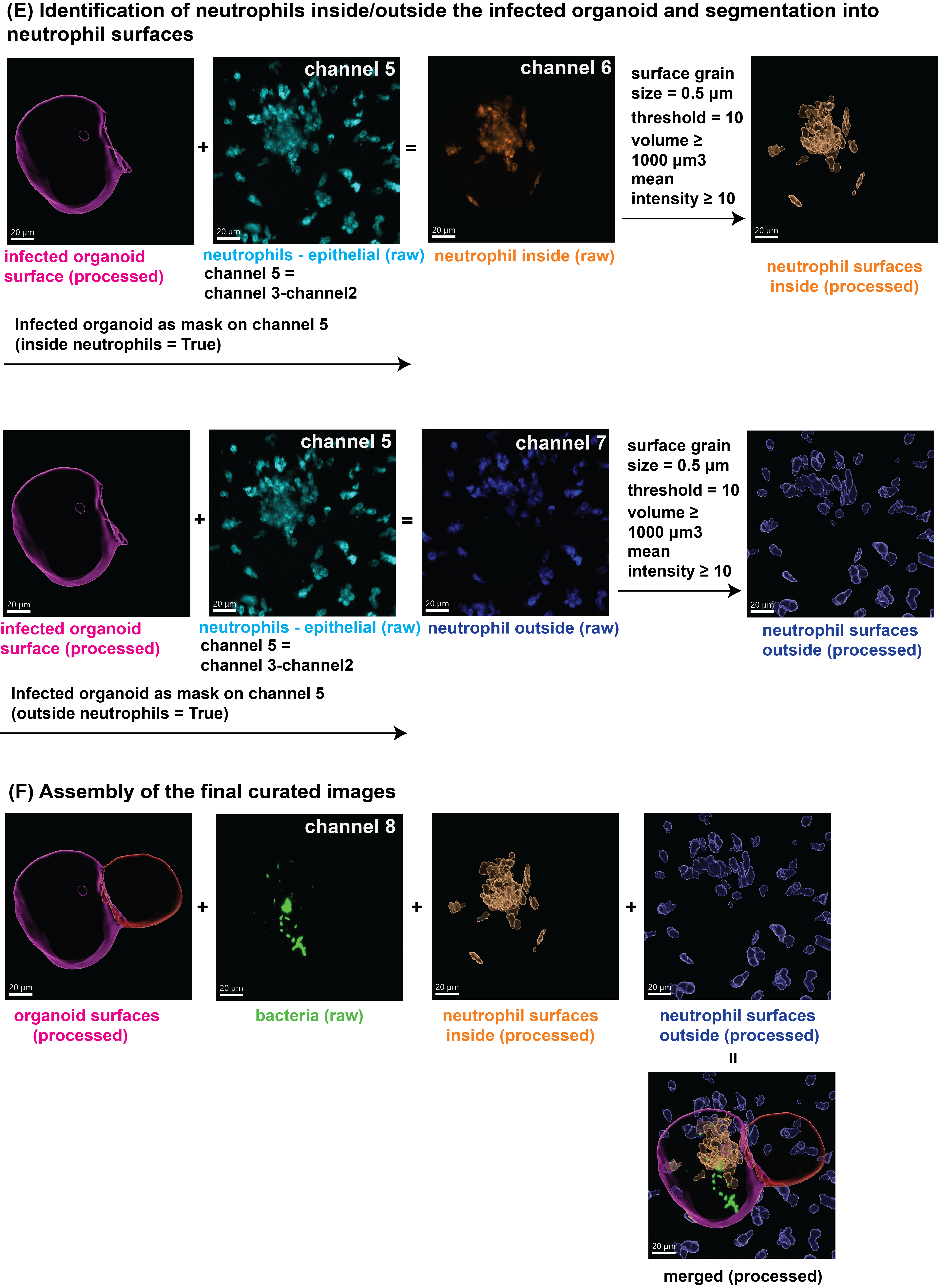

**Figure S7:** An illustrated example of the image processing pipeline for Figures 3, 4, S8 and S9.

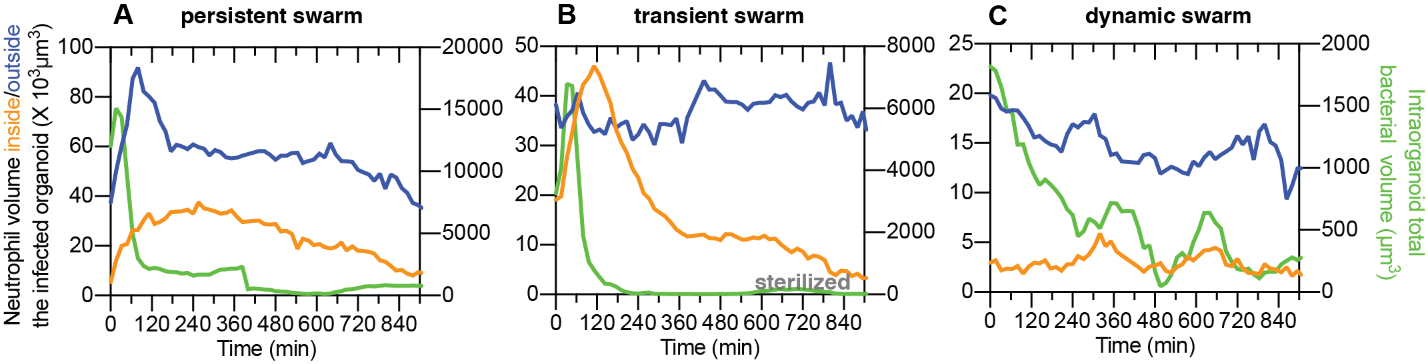

**Figure S8**. (**A-C**) Time profiles of the absolute volume of neutrophils inside the organoid, absolute volume of neutrophils outside the organoid, and the intra-organoid bacterial volume for the three swarming profiles presented in Fig. 3A-C, respectively.

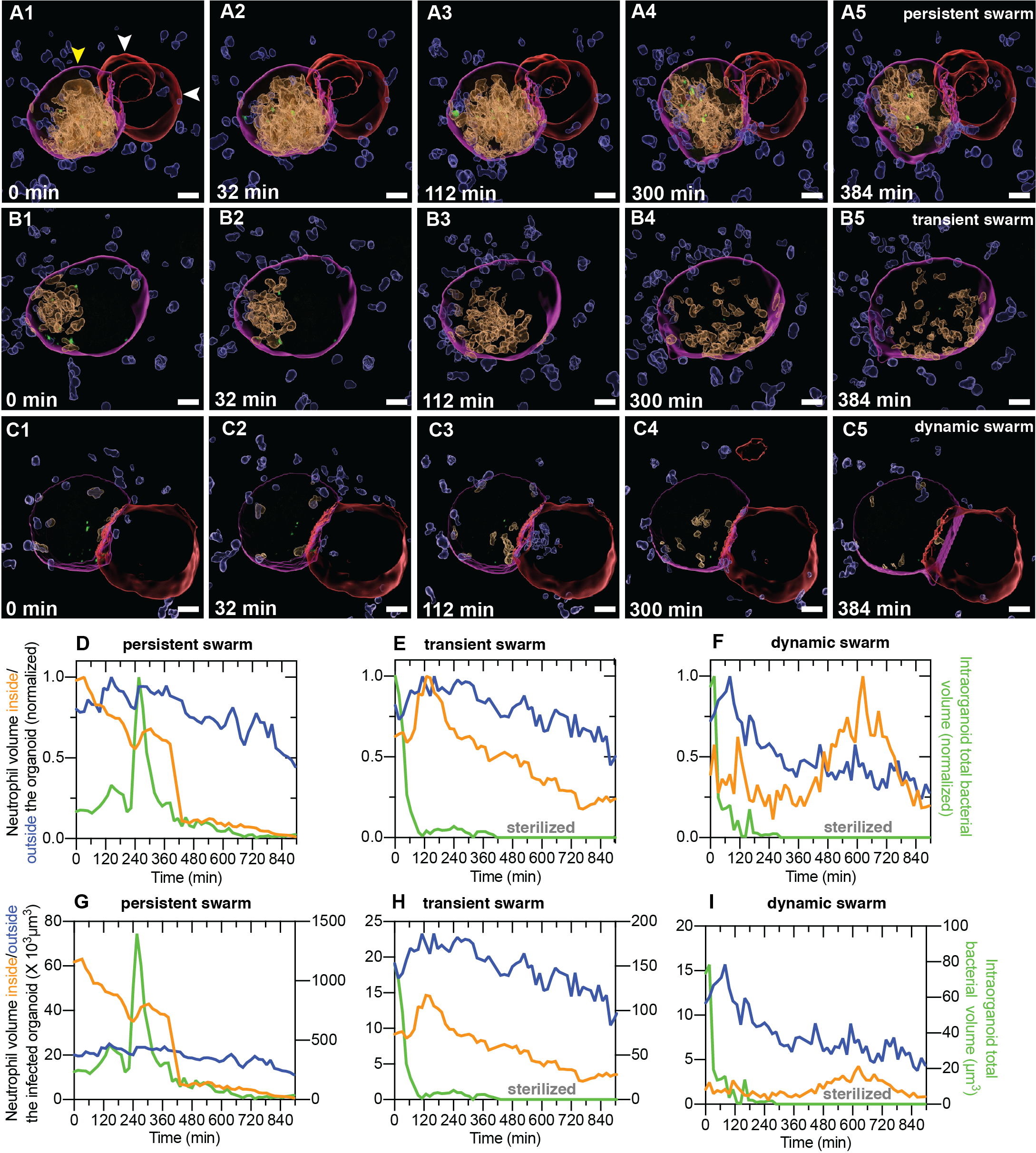

**Figure S9.** Additional examples of snapshots from time-lapse imaging showing the formation of a persistent neutrophil swarm (**A1-A5**), a transient neutrophil swarm (**B1-B5**), and a dynamic neutrophil swarm (**C1-C5**) in response to intra-organoid bacterial infection. In all panels, neutrophil surfaces (intra-organoid, amber; extra-organoid, blue) and organoid surfaces (infected, magenta; uninfected, red) were generated via an analysis pipeline using Bitplane Imaris. Bacteria (green) are shown without processing to identify individual cells. (**D-I**) Time profiles of the volume of neutrophils inside the organoid, volume of neutrophils outside the organoid, and the intra-organoid bacterial volume for the three profiles presented in (**A**-**C**), respectively. In each case, the volume normalized to the maximum volume during the course of the experiment (**D**-**F**) or the absolute volume (**G-I**) is shown.

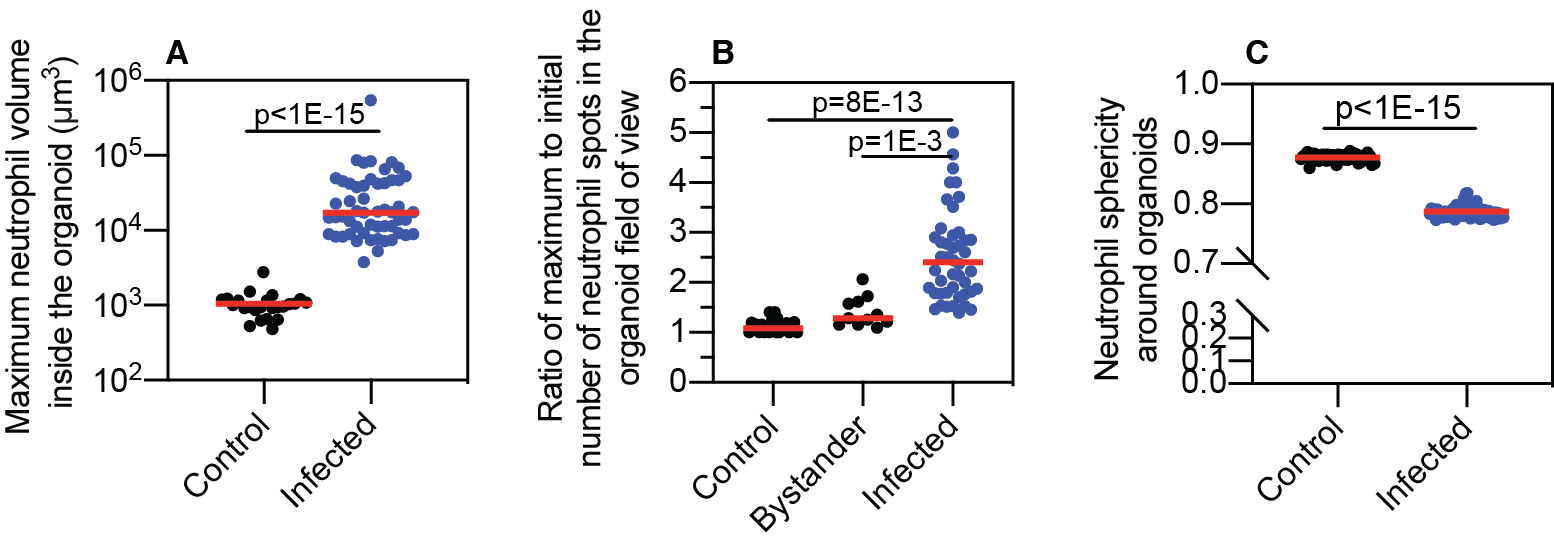

**Figure S10. (A)** Scatter plot showing the maximum neutrophil volume inside organoids during the experimental time course. Infection significantly increases neutrophil migration. P<1E-15 comparing control (n = 24) and infected (n = 54) organoids. P-values calculated using a Mann-Whitney test. (**B**) Forward migration towards organoids also increases the concentration of neutrophils surrounding the organoid for infected organoids but not for uninfected bystander organoids or control organoids. P=8E-13 for control (n = 24) vs. infected (n = 46) organoids. P=1E-3 for bystander (n = 11) vs. infected (n = 46) organoids. P-values calculated using a Kruskal-Wallis ANOVA test. (**C**) Migration can also be characterized by a reduction in the sphericity of the shape of neutrophils surrounding infected organoids. P<1E-15 for control (n = 64) vs. infected (n = 66) organoids. P-values calculated using a Mann-Whitney test.

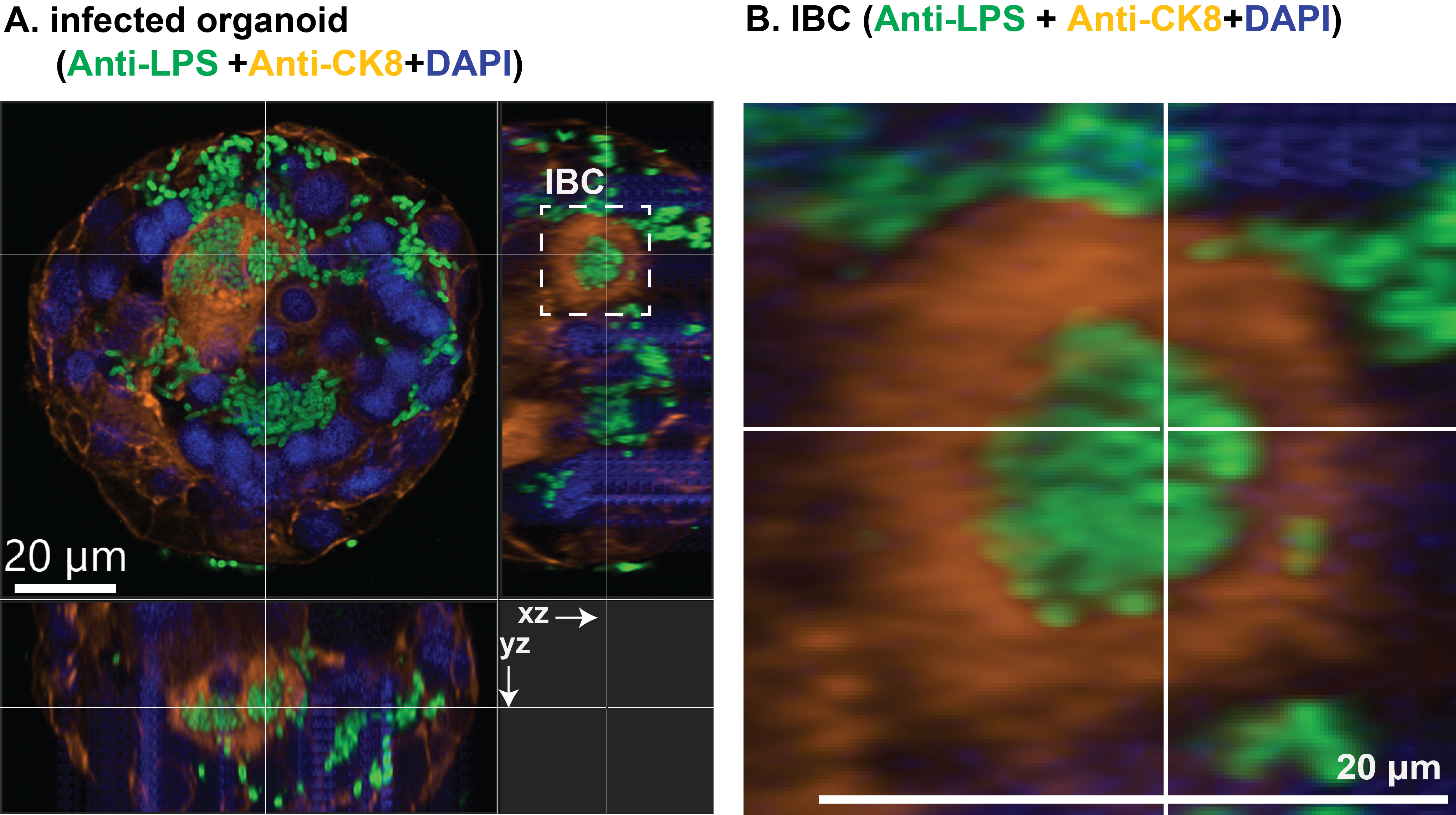

**Figure S11.** (**A**) Orthogonal sections of an infected organoid. Bacteria are identified with an anti-LPS antibody (green). An IBC within a CK8+ cell is indicated by the dashed white line. (**D**) Zoom-in to the IBC shows bacteria within a CK8+ cell. Nuclear labelling is indicated in indigo. Scale bars, 20 μm in (**A, B**).

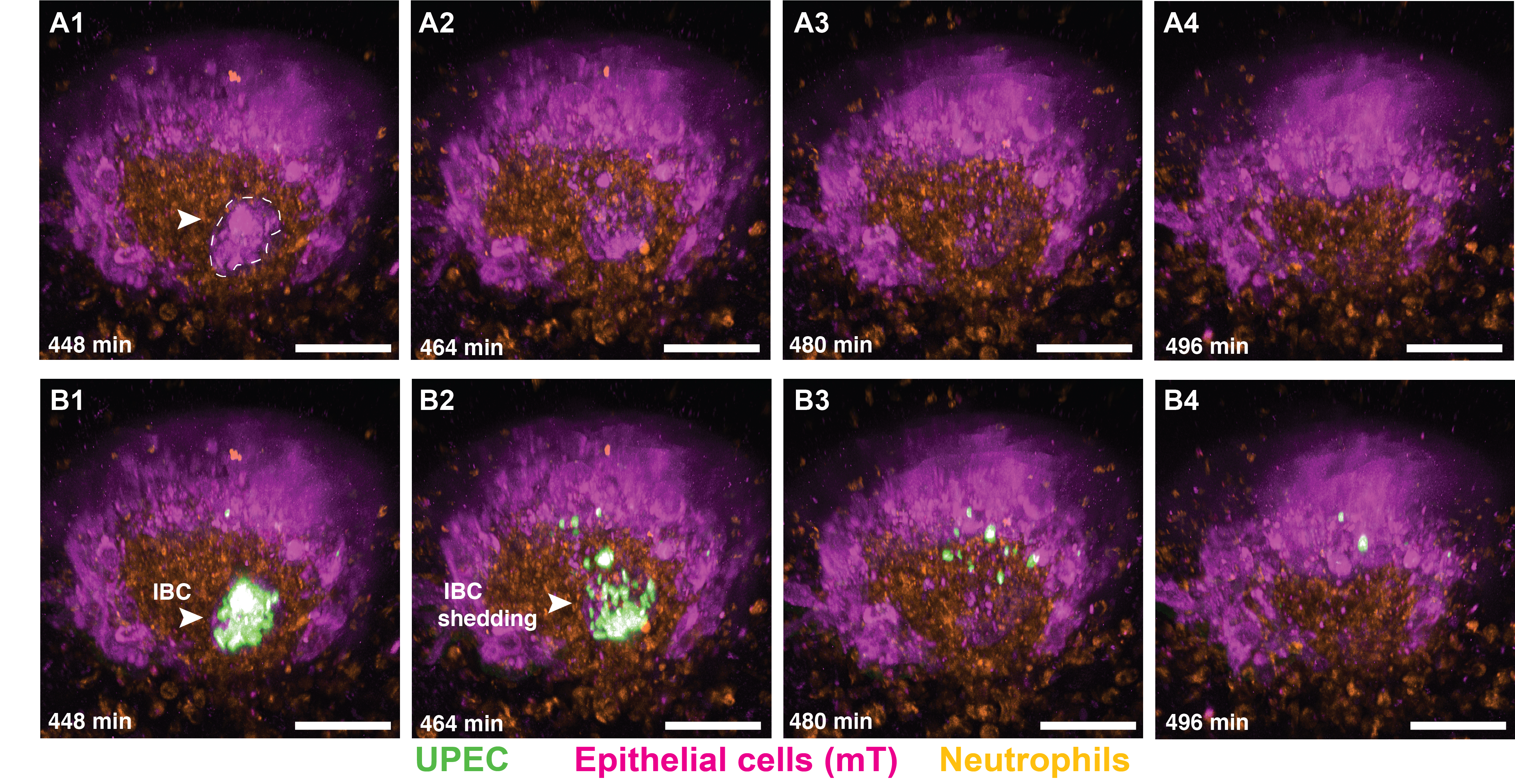

**Figure S12:** 3D views of raw data images for the time series depicted in Fig. 4, A3-A5. (**A1**-**A4**) Time series showing an overlap of the epithelial cell channel and the neutrophil channel. The dashed white line indicates the location of the IBC, which does not coincide with neutrophil staining (**B1**-**B4**). Time series showing the overlap of the epithelial, neutrophil, and bacterial channels. The intact IBC is indicated in **B1**, which is followed by shedding of the IBC and uptake of the bacteria by neutrophils (**B2**-**B4**). Scale bars, 20 μm in (**A1-A4, B1-B4**).

**Figure S13. (A)** Fluorescence image of the infected organoid used for SBEM obtained using a confocal microscope. False colouring indicates epithelial cells (grey), neutrophils (cyan), and IBC (bacteria within the IBC in yellow). (**B1-B4**) Slices from the same infected organoid acquired from SBEM, which highlight the organoid lumen (brown) shown in Fig. 5A (**B1**), neutrophils (cyan) shown in Fig. 5B, the IBC (area of IBC is shown in purple) shown in Fig. 5C (**B2**), the epithelial cell containing intracellular bacteria shown in Fig. 5D (**B3**), and the location of pericellular bacteria shown in Fig. 5E (**B4**). (**C**) Images from the Blender model of a magnified view of two neutrophils that contain bacteria. The first neutrophil (cyan) is present inside the lumen (brown) and is indicated with a red arrowhead. The second neutrophil outside the lumen is indicated with a white arrowhead. Bacteria located within the neutrophils and the lumen are colored grey and violet, respectively. Scale bars, 20 μm in (**A, B1-B4**).

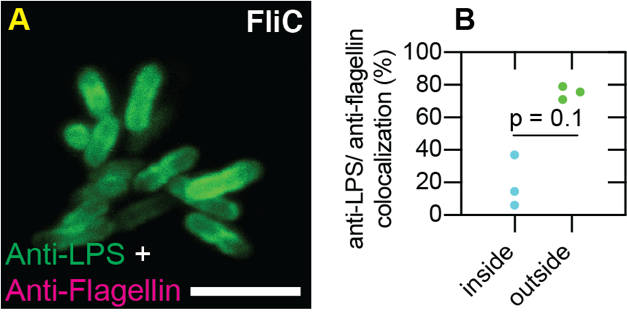

**Figure S14. (A)** UPEC Δ*fliC* mutant does not stain with anti-flagellin antibody. (**B**) Mander’s overlap coefficient **a** calculated for colocalization of anti-LPS and anti-flagellin staining on bacteria found inside and outside organoids (segmented as shown in Fig. 6A). Data are from n=3 organoids. P-values calculated using Mann-Whitney test. Scale bar, 5 μm.

### Supplementary Tables

| Expt1 | Exp2 | Exp3 |
| --- | --- | --- |
| 1155 | 680 | 345 |
| 835 | 1005 | 285 |
| 765 | 935 | 425 |
| 450 | 855 | 525 |
| 1315 | 515 | 485 |

**Table S1:** Colony forming units of UPEC within a 1 nl injection volume from three different microcapillaries. The rows of the table represent technical replicates.

| **Primer** | **Sequence (5' - 3')** |
| --- | --- |
| *Tnfa* forward | GGT GCC TAT GTC TCA GCC TCT T |
| *Tnfa* reverse | GCC ATA GAA CTG ATG AGA GGG AG |
| *Cxcl2* forward | CAT CCA GAG CTT GAG TGT GAC G |
| *Cxcl2* reverse | GGC TTC AGG GTC AAG GCA AAC T |
| *Il6* forward | TAC CAC TTC ACA AGT CGG AGG C |
| *Il6* reverse | CTG CAA GTG CAT CAT CGT TGT TC |
| *Il1b* forward | TGG ACC TTC CAG GAT GAG GAC A |
| *Il1b* reverse | GTT CAT CTC GGA GCC TGT AGT G |
| *Ccl2* forward | GCT ACA AGA GGA TCA CCA GCA G |
| *Ccl2* reverse | GTC TGG ACC CAT TCC TTC TTG G |
| *Ccl3* forward | ACT GCC TGC TGC TTC TCC TAC A |
| *Ccl3* reverse | ATG ACA CCT GGC TGG GAG CAA A |
| *Ccl4* forward | ACC CTC CCA CTT CCT GCT GTT T |
| *Ccl4* reverse | CTG TCT GCC TCT TTT GGT CAG G |
| *Il10* forward | CGG GAA GAC AAT AAC TGC ACC C |
| *Il10* reverse | CGG TTA GCA GTA TGT TGT CCA GC |

**Table S2:** Primers for qRT-PCR used in this study.
